# Recruitment of Homodimeric Proneural Factors by Conserved CAT-CAT E-Boxes Drives Major Epigenetic Reconfiguration in Cortical Neurogenesis

**DOI:** 10.1101/2023.12.29.573619

**Authors:** Xabier de Martin, Baldomero Oliva, Gabriel Santpere

## Abstract

The proneural factors of the basic-helix-loop-helix (bHLH) family of transcription factors coordinate early processes of neurogenesis and neurodifferentiation. Among them, *Neurog2* and *Neurod2* subsequently act specifying neurons of the glutamatergic lineage. The disruption of proneural factors, their target genes, and the DNA motifs they bind, have been linked to various neuropsychiatric disorders. Proneural factors operate on the DNA forming homodimers or heterodimers with other bHLH factors and binding to specific motifs called E-boxes, which are hexanucleotides of the form CANNTG, composed of two CAN half sites on opposed strands. These E-box motifs are highly enriched in regulatory elements that become active during corticogenesis. Although neurogenesis and neurodifferentiation appear to rely heavily on the activity of E-boxes, our understanding of the specific dynamics of DNA binding and partner usage throughout neurogenesis and neurodifferentiation remains largely unknown.

To shed light on this critical facet of neural development, we conducted a comprehensive analysis leveraging ChIP-seq data of NEUROG2 and NEUROD2, paired with time-matched single-cell RNA-seq and ATAC-seq assays and DNA methylation data, collected from the developing mouse brain. Our analyses revealed that distinct trajectories of chromatin accessibility are selectively linked to specific subsets of NEUROG2 and NEUROD2 binding sites and E-boxes. Notably, while E-boxes composed of CAT-CAG half sites or two CAG half sites are more commonly found within their binding sites, E-boxes consisting of two CAT half sites exhibit a striking enrichment in developmentally dynamic enhancers. These CAT-CAT E-boxes also manifest substantial DNA demethylation effects throughout the process of neurodifferentiation and display the highest levels of evolutionary constraint. Aided by a combination of a detailed DNA-footprinting and structural modeling approach, we propose a compelling model to explain the combinatorial action of bHLH factors across the various stages of neurogenesis. Finally, we hypothesize that NEUROD2 acts as a chromatin remodeler in cortical neurodifferentiation by binding CAT-CAT E-boxes as a homodimer, a mechanism that could be extended to other members of this bHLH class of transcription factors.

## INTRODUCTION

Mammalian cortical neurogenesis initiates when the self-renewing population of neural stem cells (NSCs) begins to divide asymmetrically to give rise to neurons or intermediate progenitor cells (IPCs), which in turn can go through several proliferative divisions before becoming neurons. Initially, the newborn neurons undergo morphological changes, from multipolar to bipolar, and migrate towards the cortical plate, along the basal process of the NSCs^1^. Finally, the neurons settle in the cortical plate and undergo terminal differentiation, sending their axons to distant targets, growing dendrites, and forming synapses^2–4^. These dynamic processes imply intermingled axes of cellular differentiation and maturation^5^, which are accompanied with extensive modifications of the transcriptome and epigenome. These major reconfigurations are primarily orchestrated by a subset of transcription factors (TFs)^6–8^.

It is well established that proneural transcription factors of the basic-helix-loop-helix (bHLH) family represent major early coordinators of the gene regulatory networks that drive cortical neurodevelopment^9,10^. Within this group, the expression of Neurog factors in neural progenitors is necessary and sufficient to specify a glutamatergic neuronal lineage, while simultaneously inhibiting the GABAergic lineage^10,11^, and they redundantly mediate in the acquisition of laminar identity of deep-layer neurons^12^. Neurog factors have been also implicated in neuronal migration, for instance enabling the multipolar-bipolar transition by activating the expression of *Rnd2*^13^. Furthermore, NEUROG2 has been shown to activate NeuroD genes in intermediate progenitor cells and early neurons^14,15^. This subsequent expression of NeuroD factors (*Neurod1, Neurod2 and Neurod6*) mediates the process of maturation of the newborn neuron into fully differentiated excitatory glutamatergic neurons^16^. Indeed, ChIP-seq analysis of NEUROD2 in the mouse brain followed by validation experiments indicates that NEUROD2 regulates the expression of many key TFs responsible for neuronal subtype specification (e.g., Fezf2, Bcl11b, Satb2 and Cux1)^17^, placing NeuroD factors at the root of glutamatergic neuron ontogeny.

Knockout and functional analyses revealed that NeuroD factors jointly regulate several aspects of migration and axonogenesis^18^. Single knockouts reported milder defects in neuronal migration and axonogenesis than double or triple knockouts, supporting their functional redundancy in those early processes^18,19^. The expression of these factors diverges in the mature neurons that have settled in the cortical plate: while the expression of *Neurod2* and *Neurod6* is maintained lifelong, albeit at lower levels, the expression of *Neurod1* is lost. Consistent with this later decoupling, the knockout *Neurod2* mice show more severe defects in functions characteristic of mature neurons, specifically in synaptic functions and excitability^20,21^. Interestingly, the genes regulated by *Neurod2* that mediate these functions are highly enriched in genetic variants associated with neuropsychiatric phenotypes, most prominently, autism spectrum disorder (ASD)^21^. Mutations in *NEUROD2* have been associated with a range of brain disorders, including epilepsy, ASD, and intellectual and cognitive disability^21–24^. In line with this, *Neurod2* deficient mice presented an early onset of epilepsy (Chen et al., 2016) and autistic-like behavior^21^.

Proneural factors, like most bHLHs, modulate gene expression by binding to DNA in dimeric form, either as homodimers^25–27^, by forming heterodimers with other proneural factors (e.g., NEUROG1-NEUROG2^25^, NEUROG2-ASCL1^28,29^ or NEUROG2-NEUROD4^30^), or with E-proteins, which are encoded by a more broadly expressed group of bHLH factors: *Tcf4*, *Tcf3*, and *Tcf12*^31–33^. Notably, mutations in *TCF4* are the cause of Pitt-Hopkins syndrome^32^, a neurodevelopmental disorder characterized by intellectual and motor disabilities, and have been associated with schizophrenia and autism. In addition, TCF4 targets have been also associated with schizophrenia^34,35^. Several studies have shown that experimental perturbation of *Tcf4* disrupts multiple aspects of neurodevelopment, predictably showing phenotypic overlaps with Neurog and NeuroD factors knockouts, including the pace of neurogenesis, neuronal migration, morphology and subtype specification, dendrite and synapse formation, and the establishment of interhemispheric connections^35–38^.

BHLH transcription factors bind a consensus CANNTG motif known as the E-box, being the preference for the central dinucleotide variable among the subclasses of bHLH factors. Each bHLH monomer binds one CAN half-site in opposing strands, hence we can refer to E-boxes by their 5’-3’ oriented half-sites. Heterodimers of proneural factors with E-proteins bind preferentially to CAT-CAG motifs^32,39,40^, and also to CAG-CAG and CAG-CAC motifs^39,41^. Importantly, *in vitro* studies coupled to ChIP-seq have suggested that E-box usage displayed by a given bHLH factor can imply functional differences. For instance, when NEUROD2 binds DNA through the CAG-CAG motif, these binding sites are more commonly shared with other non-neuronal bHLH factors, show reduced capacity to activate target gene expression, and these target genes are associated to general cellular processes^42^. Conversely, CAT-CAG motifs are bound more specifically by NEUROD2, the target genes associated to these motifs are more strongly transactivated and are associated with neuronal functions^42^. However, the correlation between E-box preference and neuronal developmental processes in vivo remains unknown.

In recent years, large-scale single-cell analyses have enabled the evaluation of cell type-specific developmental trajectories of chromatin accessibility in the neocortex. DNA motif enrichment analyses in these dynamic regions showed that E-boxes associated with proneural factors are the most highly enriched motifs in regions that become accessible and peak during early stages of neurogenesis^7,43–46^. Furthermore, E-boxes also present significant enrichment in regions involved in chromatin looping and that undergo CpG demethylation in neurodifferentiation, with NEUROG2 and NEUROD2 proposed as contributors to these effects^47–50^. Importantly, the numerous E-boxes residing in the accessible chromatin regions of developing neurons are enriched in mutations associated with ASD and Parkinson’s disease^44,51^.

While neurogenesis and neurodifferentiation appear to rely heavily on the activity of E-boxes, our understanding of the precise E-box utilization by the proneural bHLH factors and E-proteins remains limited. To shed light on this critical aspect of neuronal development, we conducted a comprehensive analysis leveraging ChIP-seq data of NEUROG2 and NEUROD2, in conjunction with time-matched single-cell gene expression and chromatin accessibility data obtained from the developing mouse cortex. Our analyses reveal that distinct trajectories of chromatin accessibility are selectively linked to specific subsets of NEUROG2 and NEUROD2 binding sites, with CAT-CAT E-boxes displaying a remarkable enrichment in developmentally dynamic enhancers and exhibiting the highest levels of evolutionary constraint. Finally, we harmonize observations derived from multiomics, detailed DNA-footprinting, HT-selex, and structural modeling, into a comprehensive model delineating the combinatorial mechanisms of proneural factors and E-proteins throughout neurogenesis.

## RESULTS

### Transcriptional dynamics of proneural factors and E-proteins during mouse neurogenesis

We set up to investigate the mechanisms of DNA binding and action of proneural factors during cortical neurogenesis. After an exhaustive screening of appropriate ChIP-seq datasets^52^, we identified and gathered good quality ChIP-seq data for NEUROD2 and NEUROG2^17,53^ together with single-cell transcriptomics and chromatin accessibility data^50^, all derived from mouse cortex at embryonic day 14.5 (E14.5), a period of major neurogenic activity. Prior to the ChIP-seq analysis, we characterized the expression of Neurog factors, NeuroD factors and E-proteins throughout the entire neurogenic axis, encompassing neural stem cells (NSCs), intermediate precursors cells (IPCs), and three populations of excitatory projection neurons: newborn neurons (PN1), migrating neurons (PN2), and mature neurons (PN3) (**Fig. 1A**). These data show that the expression of *Neurog2* begins in NPCs and peaks in IPCs, a pattern that is mirrored by *Neurog1*, albeit with lower expression levels. Conversely, the expression of *Neurod2* and *Neurod6* starts in IPCs but reaches its peak in immature excitatory neurons (PN1 and PN2) and it remains detected in mature neurons. In contrast, Neurod1 demonstrated later and weaker expression in immature neurons, gradually disappearing at PN2 (**Fig. 1A**). The fluctuating expression of the proneural factors contrasted to that of E-proteins, of which *Tcf4* consistently maintained the highest expression level across the entire developmental axis.

**Figure 1.**
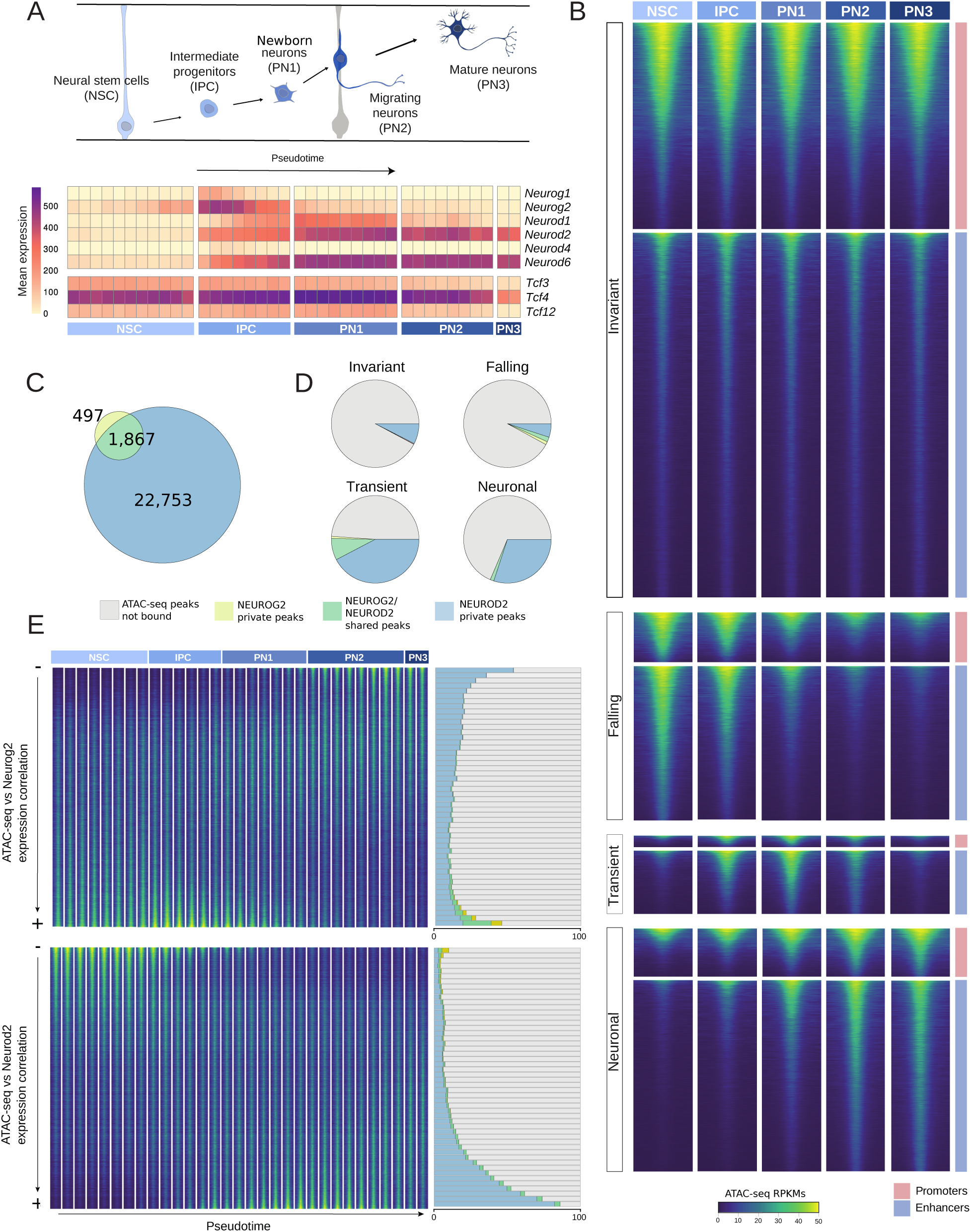
Mapping NEUROG2 and NEUROD2 binding sites to the chromatin accessibility landscape of mouse corticogenesis. (**A**) Simplified model of cortical neurogenesis, depicting the main cell types of the projection neuron lineage (top). Pseudobulk mean expression of proneural factors and E-proteins in bins of cells ordered by pseudotime (bottom). (**B**) Heatmaps depicting ATAC-seq signals measured as RPKMs in each cell-type pseudobulk, where each row represents a region of 1,500 base pairs around the summit of the ATAC-seq peaks. Regions are grouped by their main trajectories, and further subdivided by the localization within promoter or enhancer regions. (**C**) Venn diagram representing the relative amount of private and shared binding sites of NEUROG2 and NEUROD2. (**D**) Proportion of ATAC-seq peaks in each trajectory that are bound by private NEUROG2 or NEUROD2 peaks, by shared peaks, or not bound by any of them. (**E**) (Left) Heatmap representing pseudobulk signal around the summits of the ATAC-seq peaks sorted by the correlation of their accessibility with the expression of the genes. Pseudobulk samples are composed of cells grouped in increasing pseudotime bins. (Right) Proportion of ATAC-seq peaks bound by NEUROG2 and NEUROD2 in each of 50 quantiles of the correlation with the corresponding transcription factor.

### NEUROG2 and NEUROD2 binding in the chromatin accessibility landscape of mouse cortical neurodevelopment

We next aimed to position the binding of NEUROG2 and NEUROD2 within the chromatin accessibility map of cortical neurogenesis. To achieve this, we first reanalyzed ChIP-seq data for NEUROG2 and NEUROD2 in the E14.5 mouse cortex, identifying 2,356 and 26,463 peaks, respectively. The majority of NEUROG2 binding regions overlapped NEUROD2 sites, indicating that both TFs could operate on a shared subset of genomic regions (**Fig. 1C**). This smaller number of NEUROG2 peaks compared to NEUROD2 might be attributed to technical and biological reasons since at E14.5 the bulk tissue might overrepresent peaks occurring in nascent neurons, where NEUROD2 has maximum expression. Additionally, at E14.5, the neurogenic activity of NEUROG2 has been found attenuated compared to that of earlier embryonic stages^12,26^.

Secondly, in order to provide chromatin context to these binding events, we reprocessed single-cell ATAC-seq data obtained from the same developmental time and tissue^50^, obtaining four groups of peaks representing different chromatin accessibility trajectories in neurodevelopment: 1) invariant (non-variable across cell types and pseudotime), 2) falling (displaying maximum accessibility in NSCs and being subsequently closed), 3) transient (accessible in the course of neurodifferentiation and then closed in mature neurons) and, 4) neuronal (opened in the course of neurodifferentiation and still accessible in mature neurons) (**Fig. 1B**, **Extended Data Fig. 1A**). We observed that roughly half of the accessible regions were invariant, while the other half consisted mainly of regions in falling and neuronal trajectories, with only a minority in transient trajectories (**Extended Data Fig. 1B**).

The intersection of the NEUROG2 private binding sites revealed a significant enrichment in falling trajectories (Fisher’s Exact test, P=1.89e-91; OR=7.9), and to a lesser extent, transient trajectories (**Fig. 1D**, **Extended Data Fig. 1C**). Conversely, NEUROD2 private binding sites were highly enriched in neuronal and transient ATAC-seq regions (Fisher’s Exact test, P=<2.23e-308; OR=9.56, P=<2.23e-308; OR=17.16, respectively) (**Fig. 1D**, **Extended Data Fig. 1D).** The subset of NEUROG2/NEUROD2 shared binding regions was even more enriched in transiently accessible chromatin regions (Fisher’s Exact test, P=2.06e-62; OR=2.88), peaking in the period when these TFs exhibited some expression overlap (**Fig. 1D**, **Extended Data Fig. 1E).** We could observe that these highly dynamic regions enriched in NEUROD2 private and NEUROD2/NEUROG2 shared binding sites gain accessibility specifically in the lineage of excitatory projection neurons and not in other cell types present in the embryonic neocortex, including inhibitory neurons and microglial cells (**Extended Data Fig. 1F**). Moreover, they are predominantly found in enhancers compared with promoters (**Extended Data Fig. 1G-I**). Taken together, our results suggest that although NEUROG2 and NEUROD2 exhibit private binding sites indicative of specialized functions in NPCs and neurons, respectively, they may regulate a common subset of targets involved in transient functions in IPCs and early neurons.

Another inference from our analysis is that both NEUROG2 and NEUROD2 binding tend to be associated with the chromatin trajectories that resemble the most their expression patterns. With this in mind, we re-classified the ATAC-seq peaks based on the correlation of their accessibility along the neurodifferentiation axis with the expression of *Neurog2* and *Neurod2*, derived from matched transcriptomics data on the same cell populations^50^ **(Fig. 1E)**. Intersecting NEUROG2 and NEUROD2 binding sites in each of the 50 quantiles of ATAC-seq/expression correlation confirmed our previous observation: top correlated peaks exhibited the highest proportion of binding sites of the corresponding factor (**Fig. 1E**). Strikingly, 86% of the ATAC-seq peaks in the top 50th quantile most correlated with *Neurod2* expression were bound by NEUROD2, versus only 5.42% of the 1st quantile (15.87-fold enrichment). In the case of NEUROG2, although the absolute occupancy levels were lower than for NEUROD2, the difference between the top and bottom correlation quantiles was even more evident: 26.65% vs 0.11% (fold change of 242.72). We also sorted the peaks based on the correlation with the joint *Neurog2*+*Neurod2* expression, and interestingly, the binding sites shared by NEUROG2 and NEUROD2 displayed the highest enrichment in the top correlated fraction of ATAC-seq peaks (fold change of 316.06 between the first and last quantiles) (**Extended Data Fig. 1K**). Together, these results establish an association between the expression and DNA binding of NEUROG2 and NEUROD2 with a vast set of genomic regions exhibiting dynamic chromatin accessibility during neurogenesis and excitatory neuron differentiation.

### Chromatin accessibility and gene functions associated with NEUROD2 in the prenatal and postnatal neocortex

The expression of *Neurod2* subsides postnatally but is not abolished, indicating its potential role in mature neurons^21^. We set up to investigate the genomic regions targeted by NEUROD2 at E14.5 that remained targeted and/or accessible after birth. To achieve this, we utilized a NEUROD2 ChIP-seq dataset performed on the mouse cortex at postnatal day 0 (P0)^54^ and a single-cell ATAC-seq dataset of the postnatal day 56 (P56) mouse cortex^55^. While the majority of NEUROD2 peaks observed at E14.5 were undetected postnatally (**Fig. 2A**), a significant proportion of these peaks remained accessible in postnatal excitatory neurons. This finding suggests a persistent reorganization of the chromatin during neurogenesis in NEUROD2-bound regions. The subset of NEUROD2 peaks that were also detected at P0 were primarily located in accessible regions associated with transient and neuronal categories (**Fig. 2A**), highlighting a reduced set of positions that display continued *Neurod2* activity after birth.

**Figure 2.**
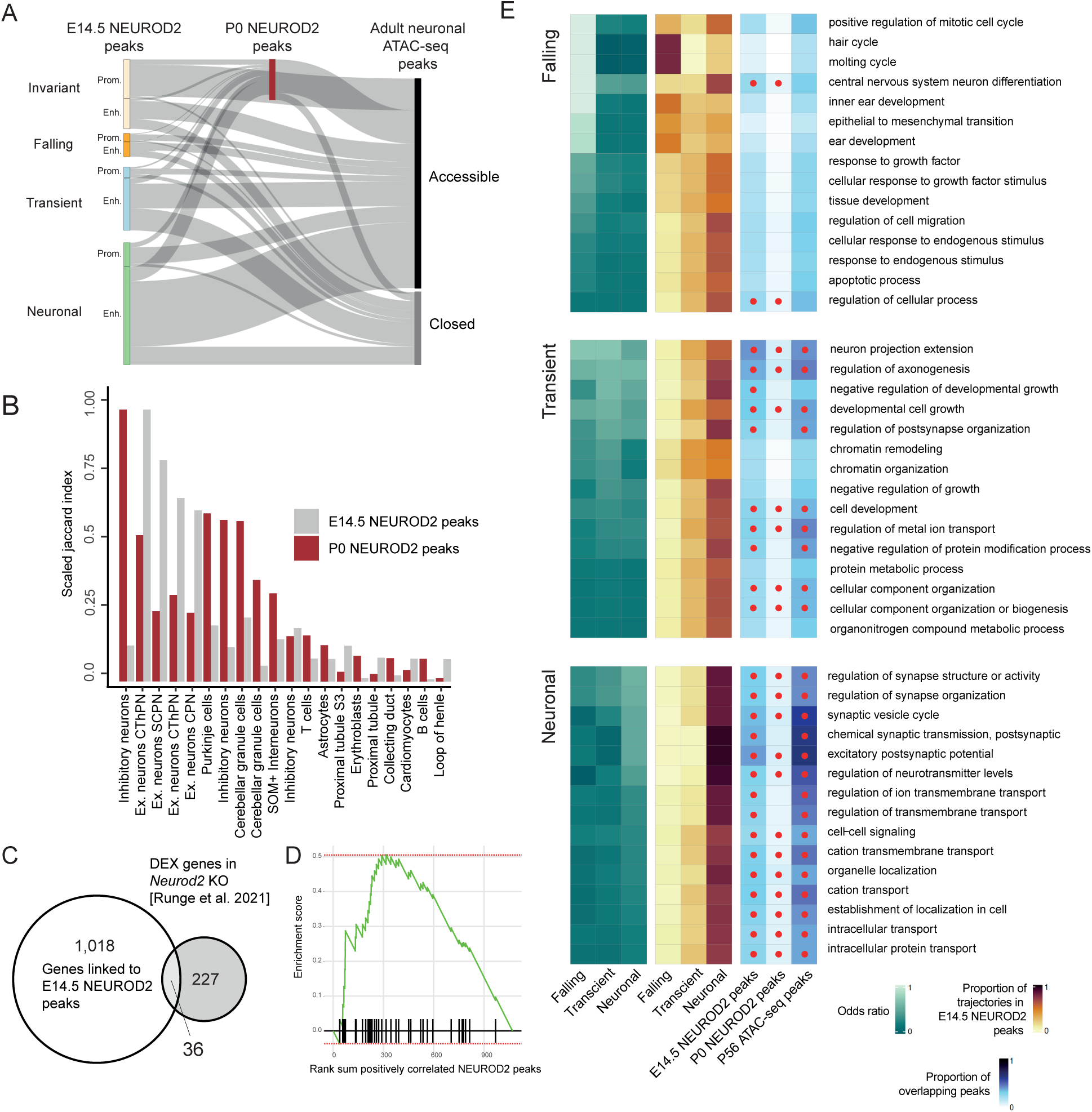
Biological functions associated with *Neurod2* in the prenatal and postnatal neocortex. (**A**) Sankey plot representing the relative number of peaks of the different trajectories which are bound or not by NEUROD2 in the P0 embryonic cortex, and that map to accessible chromatin regions in adult mouse cortical excitatory neurons. (**B**) Barplot displaying scaled Jaccard indices between NEUROD2 E14.5 and P0 peaks with regions preferentially accessible in various cell types of the mouse adult whole body. (**C**) Venn diagram depicting the intersection between genes linked with NEUROD2 peaks and genes differentially expressed upon *Neurod2* knockout in the adult mouse neocortex^21^. (**D**) Gene set enrichment analysis of the differentially expressed genes in the knockout in the adult mouse neocortex sorted by the maximum correlation of their linked peaks with *Neurod2*. (**E**) Heatmaps showing i) odd ratios of the top 15 gene ontology terms enriched in each ATAC-seq trajectory (left), ii) proportion of trajectories among the NEUROD2 peaks positively linked to genes in each ontology term (middle), and iii) fraction of peaks linked to genes in each ontology term intersecting with E14.5 and P0 NEUROD2 peaks, as well as with P56 excitatory neuron ATAC-seq peaks. Red dots indicating statistical significance, as measured by a Fisher’s exact test (right).

To explore the cell-type specificity of NEUROD2-bound accessible regions at E14.5 and P0, we leveraged a single-cell atlas of chromatin accessibility encompassing 13 adult mouse tissues^56^. Both sets of NEUROD2 peaks exhibited the highest enrichment in chromatin signatures associated with neurons (**Fig. 2B**). However, E14.5 peaks displayed maximum enrichment in elements specific to excitatory neurons, with lower overlap with peaks associated with other neuron types. In contrast, P0 peaks exhibited substantial overlap with multiple classes of neurons, including inhibitory neurons, cerebellar granule cells, and glutamatergic cortical projection neurons (**Fig. 2B**). This disparity suggests that late NEUROD2 activity regulates functions shared by diverse neuronal classes. Furthermore, we examined the overlap between NEUROD2 peaks and ATAC-seq peaks highly enriched in various subtypes of cortical excitatory neurons. Both E14.5 and P0 NEUROD2 peaks showed similar overlap among subtypes, with highest enrichment in neurons of the deepest cortical layer, L6b.

To investigate the biological functions associated with NEUROD2 peaks in prenatal and postnatal neocortex, we employed gene-enhancer links computed by Noack et al.^50^, which were based on the correlation between chromatin accessibility and gene expression within the same tissue and time point. We performed additional peak annotation strategies, including enhancer-promoter co-accessibility computed by Cicero^57^, as well as proximity to the nearest transcription start site. To evaluate the efficacy of these approaches, we quantified the proportion of inferred targets that exhibited a significant correlation with *Neurod2* expression within the subset of peaks bound by NEUROD2. Among these strategies, gene-enhancer pairs based on gene-enhancer correlation showed the highest correlation with *Neurod2* expression, prompting us to focus on this set of putative targets. Further supporting these predictions, we discovered a significant overlap between this set of targets and a group of 263 genes previously identified as differentially expressed (DEX) between the cortices of *Neurod2* KO and WT mice at P30 (Fisher’s Exact test, P=1.08x10-14; OR=5.5)^21^ (**Fig. 2D**). Moreover, the set of 263 DEX genes were found to be enriched among the top ranking *Neurod2* targets, based on their correlation values between chromatin accessibility and gene expression (Ranksum test, P= 0.046) (**Fig 2D**). Through these analyses, we compiled a list of 36 putative *Neurod2* targets (**Table 1**), some of which are known to play critical roles in cell migration and axon development (WebGestalt ORA FDR=0.0056). Notably, genes such as *Rnd2* and *Tiam2*, each associated with six NEUROD2 peaks exhibiting a maximum correlation >0.9 (**Table 1**), as well as *Nefm*, *Nrep* and members of the semaphorin-plexin signaling pathway were among the identified targets. Of particular interest is our observation regarding the gene *Rnd2,* with a critical role in neuronal migration, which was previously reported as a direct target of *Neurog2*^13^. Intriguingly, our findings at E14.5 suggest that *Neurod2* exhibits a stronger association with *Rnd2* compared to *Neurog2* (**Extended Data Fig. 2A**), which carries implications for our understanding of the roles reported for NEUROG2 in neuronal migration during late neurogenesis^13,15^. In summary, our findings highlight a subset of strongly predicted *Neurod2* targets showing sustained expression alterations in postnatal KO mice, suggesting continued regulation by *Neurod2* and/or enduring effects of the *Neurod2* KO inherited from prenatal developmental phases.

**Table 1:** Top candidate targets of NEUROD2. Maximum correlation between target gene expression and ATAC-signal is shown. Differential expression in Neurod2 KO mice from Runge et. al 2021^21^ is indicated.

| Gene | # Neurod2 peaks | Maximum correlation | log2 FC | Change |
| --- | --- | --- | --- | --- |
| Tiam2 | 6 | 0.915 | -0.494 | Down in KO |
| Nuak1 | 7 | 0.911 | -0.473 | Down in KO |
| Plxna4 | 4 | 0.911 | -0.445 | Down in KO |
| Ank3 | 3 | 0.883 | -0.313 | Down in KO |
| Tenm4 | 6 | 0.880 | -0.473 | Down in KO |
| Id2 | 3 | 0.867 | -0.916 | Down in KO |
| Abcd2 | 3 | 0.857 | -0.496 | Down in KO |
| Camk4 | 3 | 0.841 | -0.314 | Down in KO |
| Nrep | 3 | 0.839 | -0.355 | Down in KO |
| Pdzrn3 | 4 | 0.831 | -0.377 | Down in KO |
| Nefm | 3 | 0.823 | -0.410 | Down in KO |
| Smarca2 | 1 | 0.820 | -0.298 | Down in KO |
| Sema7a | 6 | 0.814 | -0.357 | Down in KO |
| Lrrc55 | 2 | 0.799 | -0.629 | Down in KO |
| 1110008P14 | 2 | 0.784 | -0.462 | Down in KO |
| Dyrk2 | 4 | 0.777 | -0.475 | Down in KO |
| Sgtb | 3 | 0.758 | -0.284 | Down in KO |
| Plch2 | 1 | 0.756 | -0.307 | Down in KO |
| Pcdh9 | 2 | 0.754 | -0.311 | Down in KO |
| Epha4 | 1 | 0.733 | -0.436 | Down in KO |
| Dgki | 1 | 0.731 | -0.397 | Down in KO |
| Tmem178b | 1 | 0.729 | -0.346 | Down in KO |
| Dock4 | 2 | 0.715 | -0.351 | Down in KO |
| Sort1 | 4 | 0.714 | -0.453 | Down in KO |
| Grin2b | 2 | 0.707 | -0.541 | Down in KO |
| Adcy1 | 1 | 0.700 | -0.461 | Down in KO |
| Rnd2 | 6 | 0.930 | 0.403 | Up in KO |
| Ly6h | 1 | 0.881 | 0.303 | Up in KO |
| Trp53inp1 | 2 | 0.880 | 0.454 | Up in KO |
| Mc4r | 2 | 0.855 | 0.618 | Up in KO |
| Sla | 4 | 0.847 | 0.645 | Up in KO |
| Pde9a | 6 | 0.836 | 0.451 | Up in KO |
| Ociad2 | 3 | 0.804 | 0.395 | Up in KO |
| Igsf21 | 3 | 0.775 | 0.376 | Up in KO |
| Ddit4 | 1 | 0.739 | 0.380 | Up in KO |
| Cdh13 | 1 | 0.724 | 0.509 | Up in KO |

To unveil the distinct functional characteristics associated with *Neurod2* targets linked to ATAC peaks of different trajectories, we conducted a comprehensive gene ontology enrichment analysis on each category. The early peaks were primarily associated with broad early developmental functions, displaying minimal overlap with *Neurod2* targets and limited association with adult excitatory neuronal ATAC-seq peaks (**Fig. 2E**). In contrast, ATAC peaks linked to transiently accessible regions exhibited significant enrichment in functions related to axonogenesis. Genes associated with these peaks displayed a high overlap with *Neurod2* putative E14.5 and P0 gene targets, as well as postnatally accessible regions. Given that previous mouse knock-out experiments have demonstrated the essential role of Neurod genes in axonal fasciculation and the development of the corpus callosum^19^, we conducted a detailed analysis of *Neurod2* targets among genes related to axonogenesis. We observed that NEUROD2 binds to regulatory elements associated with all classic axon guidance pathways, including Slit/Robo, Netrin/DCC, Netrin/Unc5, Semaphorin/Plexin and Ephrin/Eph receptors and multiple axon fasciculation molecules, such as Contactin 2 (*Cntn2*) and neurofascin (*Nfasc*) (**Extended Data Fig. 2B**). Genes linked to ATAC-peaks with neuronal trajectories produced enrichments in terms related to synaptic transmission, showing significant overlap with both E14.5 and P0 NEUROD2 peaks, as well as peaks accessible in adult excitatory excitatory neurons. Collectively, these findings indicate that NEUROD2 mediates both temporally specific and sustained regulatory activities commensurate with chromatin remodeling throughout prenatal and postnatal cortex development.

### NEUROG2 and NEUROD2 exhibit diverse E-box binding capabilities

Earlier studies have shown that variations in the central dinucleotide of the CANNTG E-box motif can affect the activity of bHLH factors, NEUROD2 among them, in different ways: it can indicate differential usage of dimerization partners, confer a bHLH factor specificity with respect to the other members of its family, and affect both binding affinity to the DNA and transactivation strength of target genes^42,52^. Consequently, we chose to explore the distribution of the types of E-boxes bound by NEUROG2 and NEUROD2 across different putative regulatory elements, stratified based on their ATAC signal versus expression correlation and trajectories of chromatin accessibility.

We first quantified the presence and centrality of E-boxes with all possible combinations of the internal dinucleotides by means of exact pattern matching. Since E-box core positions are palindromic and we disregard flanking positions, certain dinucleotide combinations are equivalent, resulting in a total count of 10 distinct types of E-boxes. It is essential to reiterate and clarify the nomenclature we employ for these E-boxes. We denote them based on the sequence in the forward strand of the two half-sites, as we believe this provides more immediate information about the individual binding of each monomer of the bHLH dimer, as we had previously proposed^52^. For instance, an E-box with the sequence CAGATG is referred to as CAT-CAG, indicating that one monomer binds a CAG half-site, while the other monomer binds a CAT half-site.

By counting and analyzing the distribution of each type of E-box around the summit of NEUROG2 and NEUROD2 peaks, we observed that CAT-CAT, CAT-CAG, CAT-CAC, CAG-CAG and CAG-CAC were enriched around peak summits (**Fig. 3A**). Conversely, the remaining five types of E-box displayed background frequencies and lacked centrality. These findings align with previous evidence showing that CAA half-sites are non-preferred by all bHLH factors^58^, and that CAC-CAC motifs are highly specific of the Myc subfamily of bHLH factors^52^ and bHLH repressors, such as Hes2, which has been found enriched in inactive enhancers during cortical neurogenesis^50^. We identified the CAT-CAG E-box as the most abundant (35.19% NEUROG2, 28.85% NEUROD2), followed by the CAG-CAG motif (21.99%, 19.76%). Although CAT-CAT motifs exhibited a lower absolute count and constituted only, 6.12%, 6.55% of all E-boxes within NEUROG2 and NEUROD2, respectively, they showed high centrality in their corresponding ChIP-seq peaks (**Fig. 3A**). CAT-CAT E-boxes displayed comparatively lower enrichment at P0 (**Extended Data Fig. 3A**), suggesting a temporally restricted role of this type of E-boxes, commensurate with *Neurod2* expression (**Fig. 1A**).

**Figure 3.**
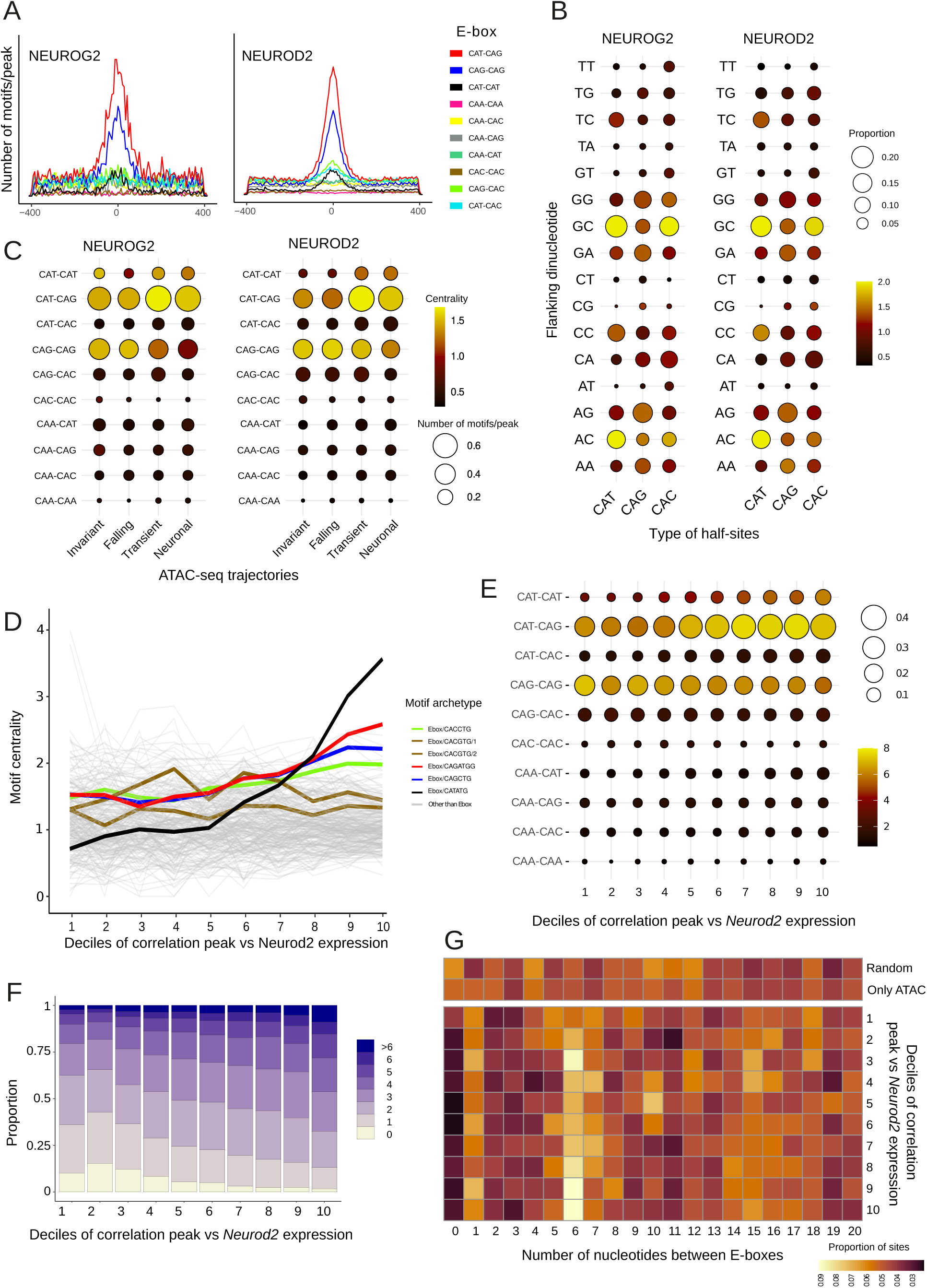
Motif grammar of proneural factors in the dynamic chromatin landscape of neurodifferentiation. (**A**) Number of E-box motifs per base pair and per peak around the summits of NEUROG2 and NEUROD2 peaks in E14.5 mouse brain. (**B**) Dot plot indicating the proportion of dinucleotides in the flanking positions of E-boxes identified in 100bp around the summit of NEUROG2 and NEUROD2 peaks, stratified for each half site (size). Color of the dots indicates the fold change of this proportion versus the corresponding proportion in motifs >200bp further away from the summit. (**C**) Dot plot representing the number of E-box per peak (100bp around the summit) in the different ATAC-seq trajectories, as well as the fold change of this number vs motifs >200bp further from the summit. (**D**) Line plot indicating motif centrality (motifs 50bp around the summit vs >250bp further) of all the archetypal motifs determined in Vierstra et al.^61^ around the summits of the ATAC-seq peaks, grouped by the correlation of their ATAC accessibility with the expression levels of *Neurod2*. (**E**) Centrality (motifs 50bp around the summit vs >150bp further) of each E-box type around the summits of the NEUROD2 peaks, binned by the correlation with *Neurod2* expression levels. (**F**) Barplot displaying the proportions of NEUROD2 peaks encompassing different amounts of E-box motifs per peak, grouped by the correlation with *Neurod2* expression levels. (**G**) Heatmap indicating the proportion of nucleotides distances between pairs of E-boxes, stratified by bins of increasing correlation between NEUROD2 peak accessibility and *Neurod2* expression levels. Spacing in random GC-content matched regions in the genome and ATAC-seq peaks with no NEUROG2/NEUROD2 peaks is also shown (top).

### E-box flanking nucleotides depend on the sequence of each half-site

We set up to identify possible biases among E-box flanking nucleotides. When we analyzed flanking regions independently in each type of E-box a clear pattern emerged (**Fig. 3B**). NEUROG2/NEUROD2-bound E-boxes containing CAT half-sites showed enrichment in GC or AC preceding dinucleotides, compared to CAG, and to a lesser extent, CAT half-sites. Previous structural studies indicate that the bHLH factors also make specific contacts with the flanking positions of the half-sites^58,59^. Thus, the half-site-specific flanking bias observed here suggests the participation of different bHLH factor dimers, with each monomer binding to its preferred half-site and associated flanking dinucleotides. Finally, while previous in vitro studies indicated increased affinity of specific bHLH factors for 5-carboxylcytosine (5caC) in the CG dinucleotide adjacent to the E-box (e.g. TCF factors)^60^, we observed no CG enrichment in the flanking positions of NEUROD2 and NEUROG2 bound E-boxes, indicating that the adjacent CpG methylation status is not a significant determinant of DNA binding affinity of these factor in vivo.

### E-boxes with CAT half-sites are preferentially enriched in chromatin regions correlated with Neurod2 expression

Each of the E-boxes examined displayed distinct enrichment patterns across the temporal trajectories of chromatin accessibility. Motifs containing a CAT half-site were more enriched in transient and neuronal trajectories, particularly within NEUROD2 peaks, in contrast to falling and invariant sites (**Fig. 3C**). Similar results were obtained by calculating motif enrichments around the summit of NEUROD2-bound ATAC-seq regions using a set of non-redundant 233 TF motif archetypes condensed by Viestra et al.^61^ (**Extended Data Fig. 3B**). Since we have previously observed that NEUROG2 and NEUROD2 occupancy in ATAC peaks highly correlated with *Neurog2* and *Neurod2* expression (**Fig 1E**), we set up to measure how E-box counts, spacing and type preference varied with the strength of the association between ATAC signal and *Neurog2* or *Neurod2* expression. We sorted all ATAC-seq peaks bound by each TF into 10 quantiles based on their ATAC signal-expression correlation. Subsequently, we calculated the centrality around the submit of the ATAC peaks using the set of 233 archetypal transcription factors motifs^61^. Notably, the motif archetypes that exhibited the most prominent association with the predicted strength of *Neurod2*-target pairs were CAT-CAT E-boxes, followed by CAT-CAG and CAG-CAG (**Fig. 3D**). When all ten E-boxes were examined through exact pattern matching around the summit of the ChIP-seq peaks of NEUROD2, the preeminence of CAT-CAT E-boxes remained as the foremost associated with peak correlation to *Neurod2* expression (**Fig. 3E**). This enrichment of CAT-CAT E-box centrality within *Neurod2*-correlated sites was evident in both enhancers and promoter regions, even more pronounced in the latter (**Extended Data Fig. 3C**). This indicates that while *Neurod2*-correlated ATAC peaks are relatively more abundant in enhancers, the subset of *Neurod2*-correlated promoters demonstrates similar properties in terms of E-box usage. Employing the same strategy with NEUROG2 private peaks yielded no association between any E-box type and *Neurog2*-peak correlation (**Extended Data Fig. 3D**), although CAT-CAT E-boxes also showed the highest centrality when ATAC-seq signal was correlated with the aggregated expression of Neurog2 and Neurod2 (**Extended Data Fig. 3E**). Lastly, the centrality of CAT-CAT E-boxes was not observed in the ATAC-seq signal in other cell types, such as interneurons, microglial and mural cells (**Extended Data Fig. 3F**), indicating that CAT-CAT E-boxes bound by NEUROD2 in the neocortex are associated with a chromatin remodeling effect specific to excitatory projection neurons. CAT-CAT E-boxes also exhibited central enrichment among the subset of ATAC-seq peaks that were correlated with NEUROD2 but not bound by NEUROD2 or NEUROG2 (**Extended Data Fig. 3G-H**). Although other Neurod factors, including the highly co-expressed *Neurod6*, could potentially explain this enrichment, the possibility of ChIP-seq false negatives in binding detection cannot be ruled out. To directly explore this possibility, we quantified the ChIP-seq signal in ATAC regions not bound by NEUROD2. We observed that ATAC regions exhibiting a high correlation with *Neurod2* displayed more NEUROD2 ChIP-seq signal (**Extended Data Fig. 3I**). This finding suggests that subthreshold NEUROD2 peaks contribute to the centrality of CAT-CAT E-boxes within ATAC-seq peaks apparently not bound by NEUROD2.

Clustering of E-boxes in regulatory elements has been suggested to enhance both binding affinity to DNA and transcriptional activation of the genetic targets, as they facilitate cooperative binding of the bHLH factors to the DNA^42,62–67^. We quantified the distribution of E-boxes in NEUROD2 peaks across the same NEUROD2-peak correlation quantiles and found that the proportion of NEUROD2 peaks with the highest number of motifs strongly correlates with these quantiles (**Fig. 3F**). We tested whether these clustered E-boxes were randomly distributed or if their spacing followed a specific pattern, as has been observed for TWIST1, which preferentially binds to 5bp-spaced paired E-boxes as a homotetramer^68^, or Ascl1 and Ascl2, which bind to closed chromatin through multiple E-boxes clustered by 10-15bp^69^. We found an over-representation of 6 nucleotide spacing between paired E-boxes within NEUROD2, but not NEUROG2, peaks, especially in NEUROD2 peaks highly correlated with ATAC-peak signal (**Fig. 3G**). This 6-bp overrepresentation is lacking in peaks displaying the lowest correlation with *Neurod2* expression, random genomic regions with similar size and nucleotide composition or ATAC peaks with no NEUROG2/NEUROD2 ChIP-seq peaks (**Fig. 3G**), suggesting intriguing functional implications.

Collectively, these findings suggest a specific grammar of NEUROD2 binding sites implicating clusters of motifs favoring certain spacing and types of E-boxes. Notably, although CAT-CAT motifs are less frequent than the other overrepresented E-boxes, namely CAT-CAG and CAG-CAG, they are by far the most enriched in highly dynamic regions in neurodifferentiation.

### CAT-CAT E-boxes are associated to larger methylation changes during neurogenesis

Previous studies have underscored the significant role played by NEUROD2 and NEUROG2 in the demethylation of CpGs in intragenic and intergenic regions associated with neuronal genes during the process of neurogenesis and neurodifferentiation. Noack et al.^48,50^ integrated whole-genome CpG methylation profiling of sorted NSCs, IPCs and PNs. with NEUROG2 and NEUROD2 ChIP-seq data of the E14.5 mouse cortex, concluding that both factors contributed to DNA demethylation throughout neuronal development. Conversely, Hahn et al.^47^ analyzed demethylated regions between PNs and NSCs and attributed the demethylation specifically to NEUROD2, which they found overlapping half of the identified demethylated regions. To reconcile these findings, we integrated the CpG methylation dataset from Noack et al.^50^ with the ATAC-seq peaks stratified based on their intersection with NEUROG2 and NEUROD2 binding sites.

This approach revealed that only those regions bound by NEUROD2 were significantly demethylated during neurodevelopment, especially from IPCs to PNs (**Fig. 4A**). NEUROG2 was found associated with CpG demethylation only when its binding sites were shared with NEUROD2, albeit at a lower magnitude, and the demethylation observed in regions only bound by NEUROG2, or not bound by NEUROG2 or NEUROD2, was minimal (**Fig. 4A**). However, regions bound by NEUROD2 exhibited varying degrees of demethylation commensurate to the initial CpG methylation levels in NSCs; while certain sites were already hypomethylated in NSC others were highly methylated. While transient and neuronal enhancers not bound by NEUROD2 also exhibited CpG demethylation, they did not reach the extent of demethylation observed in NEUROD2-bound peaks, indicating that the demethylation effects are more pronounced in the presence of NEUROD2 (**Extended Data Fig. 4A**). CpG demethylation associated with NEUROD2 sites was also linked to the correlation between *Neurod2* expression and ATAC signal (**Fig 4B**), supporting a functional link between the two events. These trends were mirrored by our measurements of chromatin accessibility, which were predictably anticorrelated with methylation levels (**Fig. 4B**, **Extended Data Fig. 4A left**).

**Figure 4.**
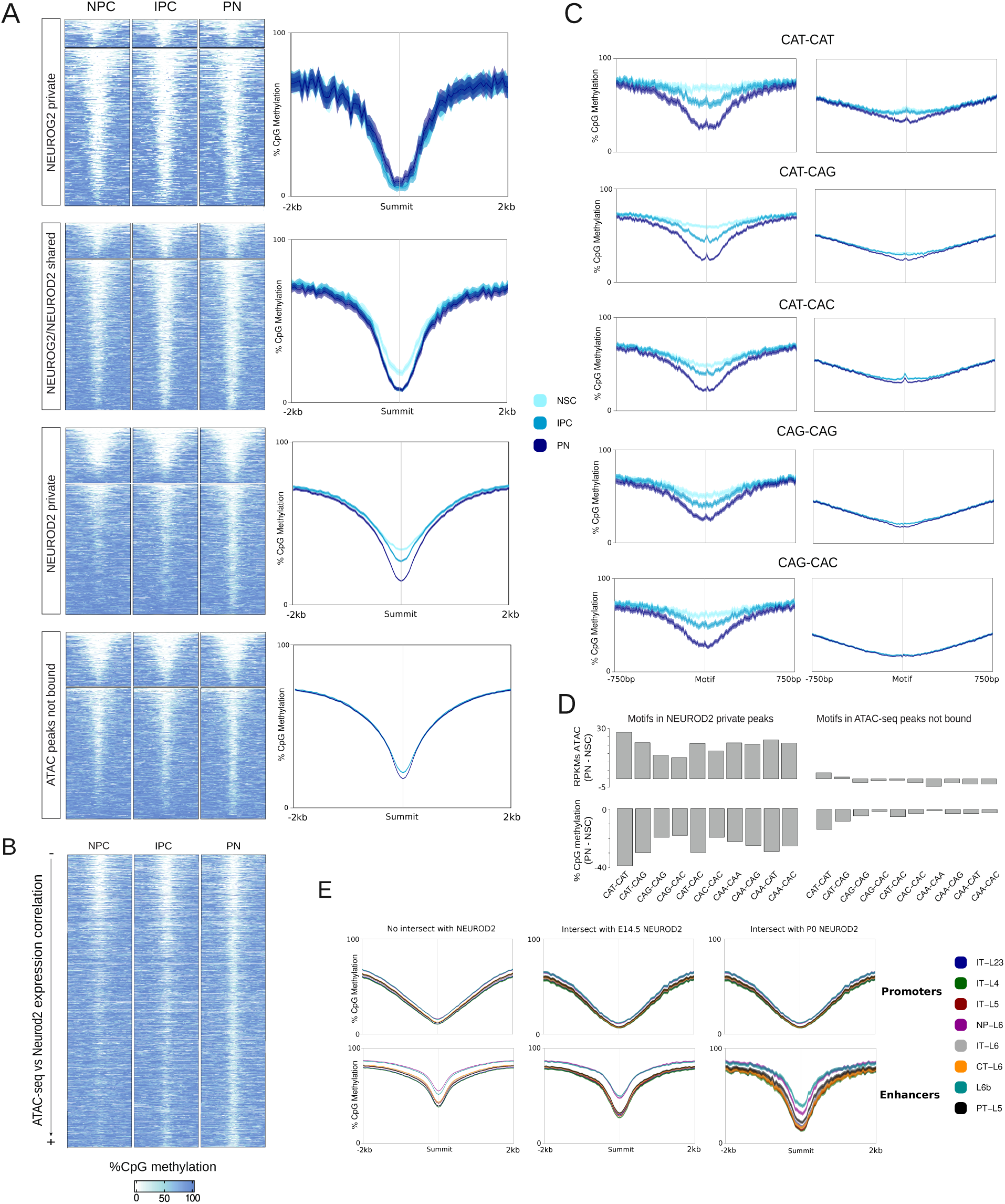
Impact of each type of E-box in the NEUROD2-associated CpGs demethylation of in the course of neurodifferentiation. (**A**) Heatmaps showing smoothed percentage of CpG methylation values in the neuronal lineage. Signal is measured in 4kb around the summits of the ATAC-seq peaks overlapping NEUROD2 or NEUROG2 private peaks, shared NEUROD2 and NEUROG2 peaks, or peaks not bound by NEUROD2 or NEUROG2 (left). Lineplot representing the average signal of the subsets of peaks on the right, with ribbons representing the standard error of the mean. (**B**) Heatmaps showing the smoothed percentages of CpG methylation around summits of ATAC-seq peaks intersecting NEUROD2 peaks sorted by the correlation of the ATAC-seq peaks with the expression levels of *Neurod2*. (**C**) Lineplots showing the average percentage of CpG methylation around different types of E-boxes within NEUROD2 peaks (left), and within ATAC-seq peaks that do not intersect with NEUROD2 (right). (**D**) Barplots representing the difference in percentage of CpG methylation measured 5bp around each type of E-box between PN and NSC (bottom). The top barplots represent the same comparisons using ATAC-seq RPKMs. (**E**) Average percentage of CpG methylation in regions overlapping NEUROD2 peaks of different cortical projection neuron cell types in the adult mouse cortex.

Next, we measured CpG methylation changes in regions surrounding each type of E-box. We found the NEUROD2 peaks containing CAT-CAT E-boxes presented the highest change in methylation levels from NSCs to PNs, followed by other CAT-containing E-boxes i.e. CAT-CAG, CAT-CAC, and CAT-CAA (**Fig. 4C-D**). All these differences were more remarkable when comparing enhancer regions versus promoter regions, which tend to be epigenetically more static (**Extended Data Fig. 4B**). Again, the CpG demethylation associated with each E-box mirrored chromatin accessibility dynamics: the motifs that were associated with a greater demethylation also were present in regions that gained more accessibility between NSC and PNs (**Fig. 4D**, **Extended Data Fig. 4B**). While E-boxes within regions not bound by NEUROD2 also reached similar methylation levels at PNs, these regions exhibited similar initial methylation levels in NSCs, resulting in a minor transition which were also observed at the level of chromatin accessibility (**Fig. 4C-D**).

Finally, to investigate the lasting stability of CpG methylation changes beyond birth, we assessed the percentage of CpG methylation in distinct subtypes of cortical excitatory neurons using single-cell methylation data from the adult mouse brain^70^. Across all subsets of peaks, we observed substantial variations in methylation levels among neuron subtypes, with intra-telencephalic types consistently displaying the lowest levels (**Fig. 4E**). Notably, both promoters and enhancers bound by NEUROD2 exhibited significantly lower CpG methylation levels in all neuron subtypes, regardless of whether we centered the analysis around the NEUROD2 ChIP-seq summit or the ATAC peak summit, in comparison to ATAC peaks not bound by either NEUROD2 or NEUROG2 (**Fig. 4E**). Interestingly, genomic regions containing NEUROD2 peaks detected at P0 displayed even lower methylation levels in adult neurons compared to those identified at embryonic E14.5 (**Fig. 4E**).

In summary, these results lend further support to the established causal relationship between NEUROD2 and CpG demethylation, adding to previous evidence^47^. Moreover, they emphasize the prominent role played by CAT-CAT E-boxes in the Neurod2-mediated reshaping of the CpG methylome during mouse corticogenesis.

### HT-SELEX uncovered a unique preference for CAT-CAT E-boxes for NEUROD2

Previous studies utilizing electrophoretic mobility shift assays (EMSA), showed that NEUROD2 and NEUROD1 bind to CAT-CAG E-boxes as heterodimers with the TCF4 factor, whereas they are not able to bind to this sequence as homodimers^32,40^. Bhlhe22, is another proneural factor exhibiting homodimerization preference in vitro and a CAT-CAT E-box enrichment in vivo^27^. This homodimer preference for CAT-CAT E-boxes extends to other non-proneural members of the same bHLH subclass, such as TWIST1^71^ or OLIG2^72^.

All this previous in vitro evidence together suggests that NEUROD2 binds to the CAT-CAT as homodimer, and to CAT-CAG as a heterodimer with E-proteins. However, it remains a plausible scenario that NEUROD2 dimers may exhibit suboptimal binding affinity to non-CAT-CAT E-boxes. High-throughput *in-vitro* assays, such as HT-selex, provide an ideal system to determine bona fide transcription factor affinities among all possible sequences tested simultaneously. Jolma et al.^73^ performed the HT-selex assay with a large number of transcription factors, including bHLH factors in their homodimeric form. They derived sequence logos indicating the consensus motif most enriched in the last rounds of the HT-selex experiment, indicating a preference of NEUROD2, NEUROG2 and the rest of proneural factors for the CAT-CAT E-box^52,73,74^. However, in these analyses the relative enrichment of each type of E-box was not explicitly investigated. To that aim, we reanalyzed the successive HT-selex rounds obtained for NEUROD2 and NEUROD2 and measured the proportion of sequences containing each type of E-box in each round. This analysis establishes the unique enrichment of CAT-CAT E-boxes through HT-SELEX (**Fig. 5A**), with no analogous secondary enrichment observed in any other E-box type, including CAT-CAG. These findings support the premise that CAT-CAT and CAT-CAG E-boxes, when occupied by NEUROD2 or NEUROG2, are operated in different partner configurations.

**Figure 5.**
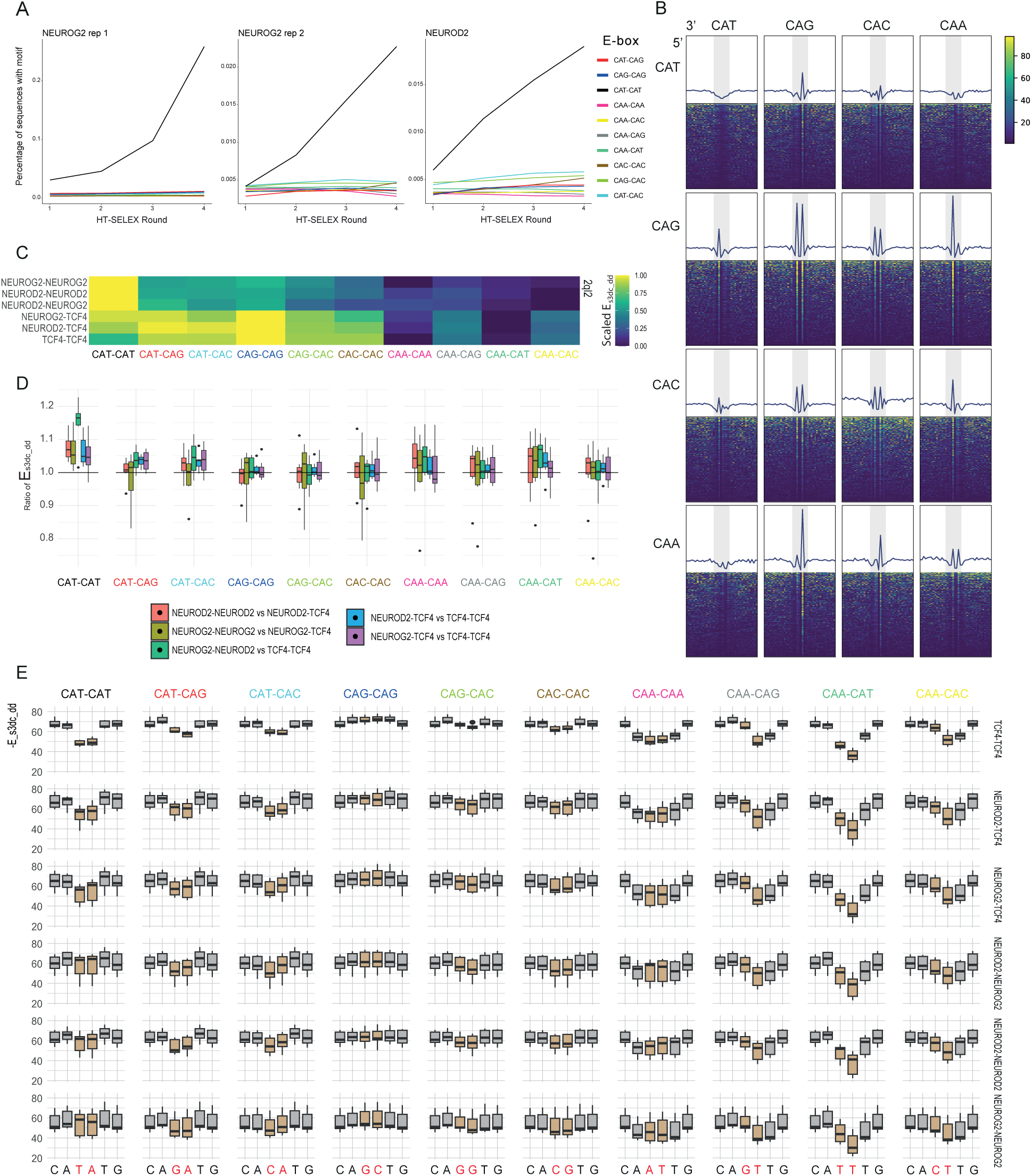
Comparison of TF-DNA binding affinity among types of E-boxes. (**A**) Lineplot representing the fraction of sequences that contain each type of E-box in the successive rounds of HT-SELEX. (**B**) Footprinting analysis showing HINT-normalized transposase cut frequency in aggregated (lineplot, up), or in individual peaks (heatmap, down), around all possible CANNTG hexanucleotides in regions bound by NEUROD2 (**C**) ModCRE calculated binding energies between various combinations of NEUROD2, NEUROG2 and TCF4 dimers in each type of E-box using the Protein Data Bank (PDB) model 2ql2 as template. Heatmap displays row-scaled normalized binding energies. (**D**) Boxplot plot showing the distribution across different PDB structures templates of the normalized binding energies for various combinations of NEUROD2, NEUROG2 and TCF4 dimers in each type of E-box. (**E**) Boxpot showing the distribution of binding affinities at the nucleotide level for each dimeric composition and E-box, using multiple PDB structures as template.

### CAT half sites carry a distinct footprinting signature

We reasoned that if dimer composition could explain differences in E-box usage, this also might lead to different footprinting signatures for each E-box type. To explore this, we conducted an E-box footprinting analysis using scATAC-seq signal overlapping E-boxes bound by NEUROG2 and NEUROD2. Transposase cleavage frequencies were computed and corrected for expected cleavage bias with HINT-ATAC^75^. When we aggregated the corrected cleavage frequency around the various E-box types within the NEUROD2 peaks, we discovered distinct footprinting signatures (**Fig. 5B**). Notably, only the CAT-CAT motif exhibited a symmetrical drop in the cleavage frequency, while the other half-site combinations displayed a peak of signal at the center of the motifs, irrespective of the strand orientation. Such spikes have been also observed in footprinting analysis of other bHLH factors, for example in CAC-CAC E-boxes bound by USF1 and have been associated with lower affinity contacts with the internal dinucleotides allowing the insertion of the transposase^76^. Specifically, we observed such signal spikes in the CAG and CAC half-sites, whereas the drop in the insertion frequency appeared in the CAT half site, implying enhanced affinity.

### Prediction of differential TF-binding affinity among E-Boxes and bHLH combinations through homology-based modeling

Interestingly, the crystal structure that was resolved by Longo et. al^59^ for the NEUROD1-TCF3 dimer binding to the CAT-CAG motif aligned with our predictions based on this reasoning and the footprinting patterns. In that structure, NEUROD1 contacts the entire phosphate backbone of its CAT half-site, while TCF3 does not contact the phosphate between the “A” and the “G” of its CAG half-site. This structural insight further supported our hypothesis regarding the role of distinct dimer compositions in influencing E-box usage. To delve deeper into the binding compatibility of different combinations of proneural factors and co-exiting E-proteins during neurogenesis, we employed a recent structure homology modeling approach called ModCRE^77^. Using this method, we systematically investigated the TF-DNA binding affinities for each type of E-box in various bHLH homo and heterodimers. ModCRE revealed higher protein-DNA binding affinities for CAT-CAT E-boxes in NEUROD2 and NEUROG2 homodimers (or NEUROD2-NEUROG2 heterodimers), and for CAT-CAG E-boxes in TCF4 homodimers or NEUROG2/NEUROD2-TCF4 heterodimers (**Fig. 5C**). E-boxes containing CAA half-sites exhibited the lowest binding affinities regardless of the accompanying half-site. The observation of a higher affinity for CAT-CAT E-boxes of proneural homodimers compared with dimers containing E-proteins, or proneural/E-protein heterodimers compared with E-protein homodimers, remained largely consistent across all seven template structures used for the initial modeling of the different dimers (**Fig. 5D**). We also quantified TF-DNA binding affinity at the nucleotide level, within each E-box. NEUROD2 and NEUROG2 homodimers, as well as NEUROG2-NEUROD2 heterodimers, exhibited the highest affinity for the internal dinucleotides of the CAT-CAT E-boxes when compared to TCF4-containing structures (**Fig. 5E**). However, when comparing different E-boxes, these proneural homodimers also showed similar affinities with the internal dinucleotide of CAG-CAG E-boxes, in contrast to the relatively lower affinity observed in the global structure compared to CAT-CAT E-boxes (**Fig. 5C-D**). Nonetheless, the nucleotide affinities were lower in absolute terms when compared to those exhibited by TCF4. TCF4 homodimers displayed a higher affinity for internal dinucleotides of CAG-CAG E-boxes, and a lower affinity in CAT-CAT E-boxes (**Fig. 5E**). In CAT-CAG E-boxes, TCF4 homodimers showed a higher affinity for the G nucleotide, as compared to the T nucleotide (**Fig. 5E**). Finally, we explored interaction differences between residues within the basic DNA-binding domain of each TF and CAT-CAT or CAG-CAG E-boxes. While most residues showed similar interaction scores with both sequences, differences surfaced in the last amino acid of the 13 residues constituting the basic domain, a defining variable position in the bHLH class^52,78^. Specifically, the Val580 of TCF4 displayed better interaction with CAG-CAG, whereas Met135 or Met125, present in the corresponding position within the sequence of NEUROD2 and NEUROG2, respectively, interacted with higher score with the CAT-CAT E-box sequence (**Extended Data Fig. 5**).

Taken together, the reanalysis of our in vitro binding assays, footprinting analysis, and structural prediction collectively suggest a univocal relationship between E-box usage and dimeric composition of proneural factors and E-proteins.

### CAT-CAT E-boxes exhibit the highest level of sequence conservation

We next investigated the level of sequence conservation within the entire set of E-boxes bound by NEUROD2 and NEUROG2. We conducted measurements of divergence by counting substitutions against the rat genome. In parallel, we assessed the sequence diversity within these E-boxes by harnessing a single nucleotide polymorphism (SNP) panel derived from 154 whole-genomes obtained from wild mice^79^. We noticed a clear relationship between both divergence and diversity and the distance of E-boxes from the peak summit (**Fig 6A** and **6B**, respectively). We then compared divergence and diversity values between those E-boxes proximal to the peak summit (50 bp around the summit), that we called Putatively Selected Sites (PSS), and E-boxes in regions distal to the summit (200-400 bp), which we called Non-Selected Sites (NSS). Our analysis of the PSS/NSS ratios revealed that CAT-CAT E-boxes display the most pronounced change, followed by CAT-CAG E-boxes, a trend observed both in terms of divergence and diversity (**Fig 6C,D**). In general, observations were consistent between NEUROD2 and NEUROG2 in terms of divergence, albeit the higher variation levels in the later due to the smaller number of NEUROG2 peaks analyzed and variants overlapping those peaks (**Extended Data Fig. 6A-B**). We contrasted diversity and divergence in PSS and NSS under the Macdonald-Kreitman framework (**Fig 6E**) using random subsets of E-boxes of the same type. Except for CAA-CAA and CAA-CAC E-boxes, all types reside in the quadrant denoting purifying selection (i.e. pPSS/pNSS < 1 and dPSS/dNSS < 1).

**Figure 6.**
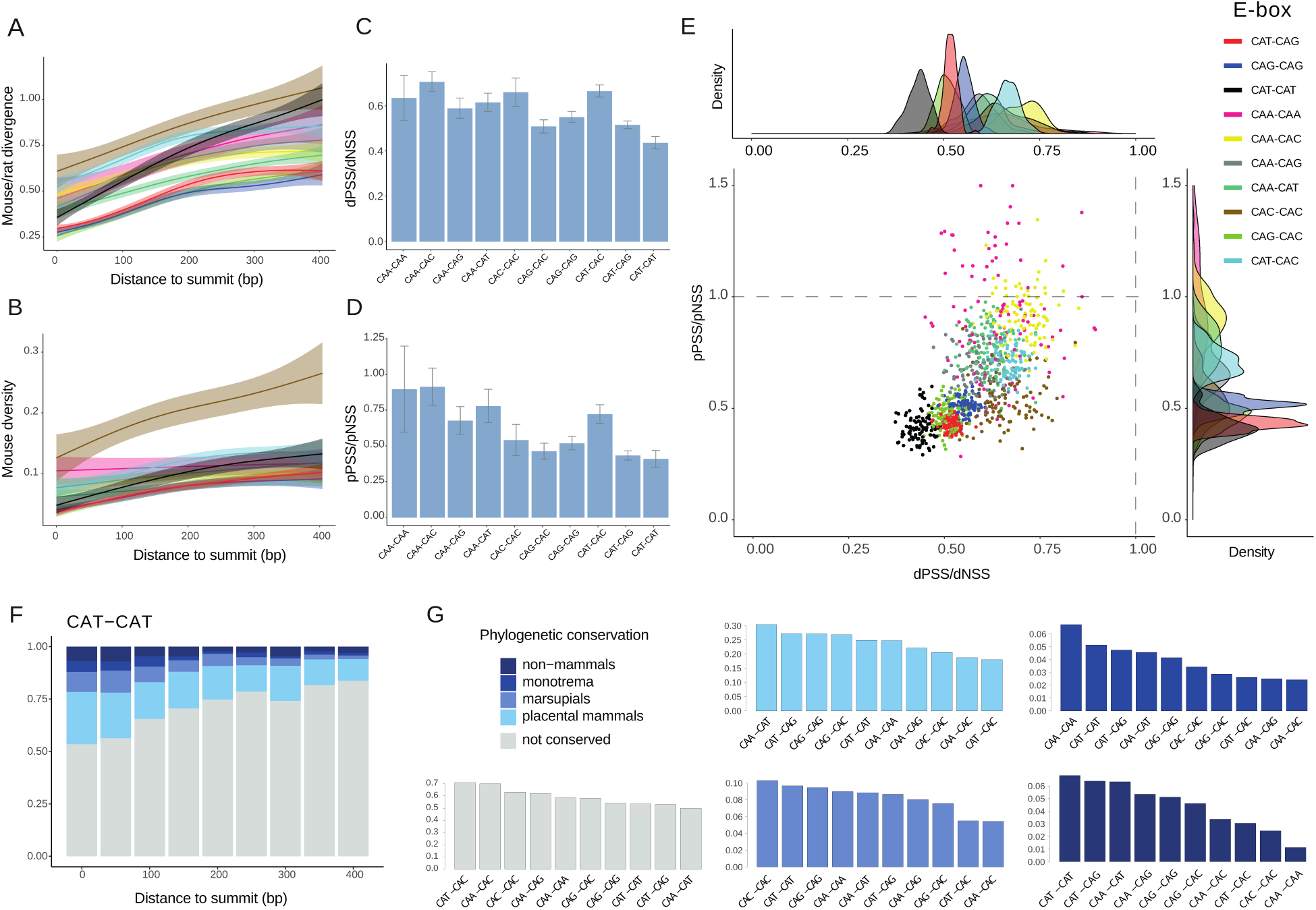
Sequence conservation of E-box motifs in NEUROD2 peaks. (**A**) Line plots showing number of SNPs within E-boxes in bins of increasing distance to NEUROD2 peak summit. (**B**) Line plots displaying the number of mouse-rat substitutions within E-boxes in bins of increasing distance to NEUROD2 peak summits. (**C**) Barplot showing the ratio of the pPSS and pNSS in each E-box type in NEUROD2 peaks. (**D**) Barplot showing the ratio of the dPSS and dNSS in each E-box type in NEUROD2 peaks. (**E**) Scatter plot showing pPSS/pNSS and dPSS/dNSS ratios in subsets of E-boxes of each type found in the summit of NEUROD2 peaks. (**F**) Barplot representing the proportion of CAT-CAT E-boxes displaying various degrees of phylogenetic conservation at bins of increasing distance to NEUROD2 peak summits. (**G**) Barplots showing the proportion of E-boxes of each type that is conserved at various phylogenetic levels.

These analyses identified CAT-CAT E-boxes as the most highly constrained motif within contemporary mice populations and extending back to the time of the split from the rat. To further explore the depth of sequence conservation among different E-box types, we used MULTIZ alignments of 60 species of vertebrates and determined the emergence of each E-box within the species tree (see Methods). Once again, we observed a clear relationship between the depth of phylogenetic conservation and the proximity to the NEUROD2 peak summit (**Fig 6F**, **Extended Data Fig. 6C**). Focusing specifically on E-boxes situated near the summit (100 bp around the summit), we found that CAT-CAT E-boxes were the most conserved across vertebrates, extending beyond mammals, and the second most highly conserved until the time of the split with monotremata (i.e. platypus) (**Fig 6G**). Following closely were other CAT-containing E-boxes, such as CAT-CAG and CAT-CAA, which also exhibited considerable levels of conservation (**Fig 6G**).

In summary, our results indicate that, although CAT-CAT E-boxes constitute only a limited portion of the overall E-box binding repertoire for NEUROD2 and NEUROG2, they display the most pronounced evolutionary constraints, as evidenced by measures of diversity, divergence, and phylogenetic depth.

### Epigenetic reconfiguration is driven by homodimeric proneural factors

One plausible explanation for the observed temporal shift between the enrichment of CAT-CAT and CAT-CAG motifs, along with the favored centrality of the former within ATAC regions, could be attributed to the relative abundance of NEUROG2/NEUROD2 and their partners with a CAG preference, namely, the E-proteins. The concentration of these factors is likely to play a pivotal role in determining the specific dimer composition that forms during neural differentiation. Notably, E-proteins are expressed at significant levels by the time *Neurod2* starts to be expressed (**Fig. 1A**). Furthermore, heterodimers between proneural factors and E-proteins seem to be more stable and exhibit longer half-lifes (e.g. 5h compared to 30 minutes of NEUROG2/E-protein heterodimers and NEUROG2 homodimers, respectively)^26,80^. According to this model, which we represent in **Figure 7**, NEUROD2/E-protein heterodimers with a CAT-CAG preference would dominate during the early stages of neural differentiation, until the increasing concentration of NEUROD2 eventually enables a greater propensity for NEUROD2 homodimerization and binding to their preferred CAT-CAT motif. These E-boxes, although relatively underrepresented compared with CAT-CAG and CAG-CAG E-boxes, display higher chromatin reconfiguration strength and demethylation effects, and are found under stronger purifying selection. The model establishes that precisely regulated relative levels of E-proteins and proneural factors are necessary to cover their binding landscape and associated targets, encompassing at least five types of centrally enriched E-boxes (**Fig. 3A**). The interaction within one cell of proneural factors, E-proteins, and their inhibitors, the ID proteins, collectively determine their DNA interaction and targets, as observed in studies of co-expression of *Neurog2* and *Tcf4*, and *Neurod1* and *Tcf12*^33,39^.

**Figure 7.**
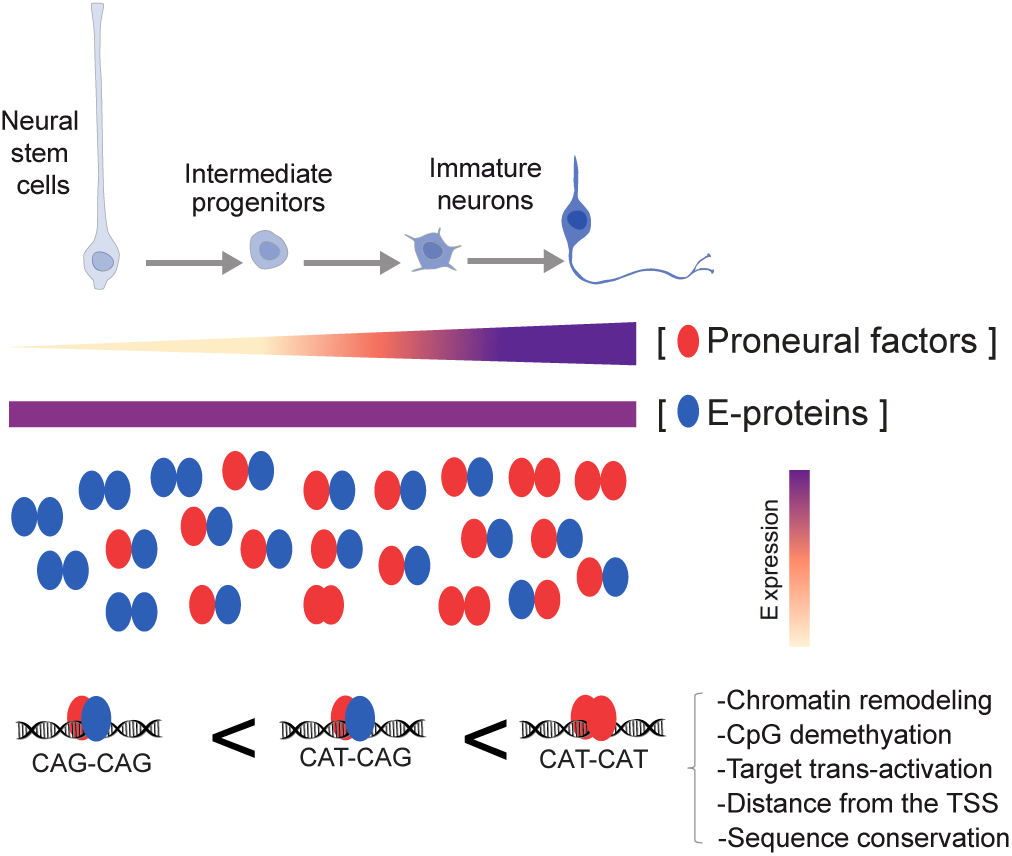
Schematics representing the properties of CAT-CAT E-boxes and the relationship between CAT-CAT occupancy and the homodimeric or heterodimeric action of proneural factors as a function of their relative concentration within the nucleus.

This model can help to reconcile the two apparently contradictory observations indicating that both the induction and repression of E-proteins results in reduced cortical neurogenesis^35,39^. We propose that while elevated concentration of E-proteins dampen the formation of proneural homodimers, crucial for binding CAT-CAT motifs in region requiring chromatin remodeling, a low concentration of E-proteins compromise the formation of proneural/E-protein heterodimers, hindering the ability of proneural factors to act upon the abundant accessible CAT-CAG and CAG-CAG E-boxes.

An illustrative example of the explanatory power of our model is provided by the study of the promoters of *Neurod1*, induced at early stages of neurodifferentiation, and Delta (Dll1), which regulates the NPC state maintenance in surrounding cells. It has been demonstrated that the induction of E-proteins enhances the ability of NEUROG2 to activate the expression of the Delta gene^26^, while inhibiting both the ability of Neurog1/2 to activate the *Neurod1* gene and their binding strength to its promoter^26,39^. By studying the E-box content and proneural binding of its promoters (**Extended Data Fig. 7A**) we can observe that the Delta promoter, which is highly accessible in progenitor cells, contains 4 CAA-CAT and a single CAT-CAG motif overlapping NEUROG2/NEUROD2 peaks, while the proneural binding sites on the Neurod1 promoter, which is remodeled to gain chromatin accessibility during neurodifferentiation, encompass a CAT-CAT, a CAT-CAG, a CAT-CAC and a CAC-CAC E-boxes. These observations made on these two well-studied promoters aligns with our model wherein the higher DNA binding affinity of proneural homodimers is required for DNA remodeling, whereby the proneural/E-protein heterodimers bind with lower affinity to regions already accessible.

## Discussion

This work establishes that CAT-CAT E-boxes bound by NEUROD2 are critical in the reconfiguration of chromatin accessibility that underpins neuronal differentiation. We reveal that NEUROD2 binds at the center of a majority of genomic regions whose accessibility and methylation levels exhibits a strong correlation with *Neurod2* expression, suggesting a previously unrecognized causative role of NEUROD2 in chromatin remodeling. During cortical neurogenesis, this major reconfiguration mediated by CAT-CAT E-boxes is highly specific to excitatory neurons. Furthermore, these CAT-CAT sites are specially conserved across evolution, as evidenced from genetic diversity, divergence rates and phylogenetic depth.

Our re-examination of HT-SELEX data, and footprinting analysis coupled with structural modeling, provides compelling evidence that homodimers of NEUROG2 and NEUROD2 exhibit a marked preference for binding to CAT-CAT E-boxes, demonstrating both selectivity and enhanced binding affinity, whereas CAT-CAG E-boxes bind to heterodimers formed in conjunction with E-proteins. Thus, by connecting observations derived from the reanalysis of functional genomics and in vitro datasets, complemented with structural analyses, we hypothesize that the variation in E-box usage reflects different proportions of homodimeric and heterodimeric forms of the Neurod factors and E-proteins, with the Neurod homodimeric form gaining maximum influence in regions gaining accessibility in immature neurons.

Our approach, which involves classifying binding sites based on the correlation between ATAC signal and TF expression, has proven to be an effective strategy for extracting functional insights from TF ChIP-seq data. Notably, NEUROD2 occupancy exhibited a strong positive correlation with this measure, as did other grammatical features of the motifs, such as clustering and spacing. This approach holds promise for broader application across other TF and cell types, enabling the identification of key regulators of chromatin reconfiguration across various lineages, and their underlying DNA-binding mechanisms.

The stratification of NEUROD2 peaks in E-boxes and ATAC trajectories clarifies the interpretation of previous studies, based on convoluted E-box enrichment in proneural factors, assigning biological roles to different types of E-boxes. For example, Fong et al. found that CAT-CAG motifs exhibit larger transactivation of NEUROD2 targets in P19 cells. In addition, targets regulated by these E-boxes were found more associated to neuronal functions, all in comparison to the CAG-CAG motifs^42^. Our analysis helps explain this observation, as we found CAT-CAG motifs to be more enriched in dynamic enhancers that become accessible during neuronal differentiation. Fong et al. interpreted that, by binding to CAT-CAG motifs, NEUROD2 acquired a certain 3D structure favoring the recruitment of coactivators of the expression. In addition, Hahn et al. suggested that by binding to CAT-CAT and CAT-CAG motifs, NEUROD2 acquired a conformation that facilitated the recruitment of the demethylating TET2 enzyme^47^. We suggest, instead, that CAT-CAT and, to a lesser degree, CAT-CAG motifs, along with motif clustering, imply a higher affinity of interaction with the DNA, which allows both binding to nucleosome-rich regions, and a longer dwell-time in the DNA, that permits the recruitment of cofactors like the TET2 enzyme and the transcriptional machinery. Those regions would become accessible upon NEUROD2 binding and subsequently demethylated, resulting in downstream genes activation. Consequently, the effects on chromatin remodeling and CpG demethylation appear comparatively reduced in lower affinity CAG-CAG E-boxes, which are more prevalent in regions which we have already identified as more accessible and hypomethylated in neuronal progenitors.

Importantly, the centrality of CAT-containing E-boxes within ATAC-seq peaks, particularly those marking dynamic chromatin regions during neurogenesis and neurodifefrentiation, strongly suggest a causal role for these E-boxes in chromatin remodeling. Furthermore, the high occupancy of these E-boxes by NEUROD2, along with the observed correlation between ATAC signal and *Neurod2* expression, implies a direct involvement of NEUROD2 in this remodeling process. It is worth noting that NEUROD1, a bHLH factor with a recognized pioneer capacity^81^, and *Neurod6*, are largely co-expressed with *Neurod2* and could also potentially bind to the same E-boxes. This could explain why some centrally enriched E-boxes in transient chromatin regions do not show NEUROD2 binding in our dataset, even though we demonstrated the contribution of false negatives from NEUROD2 ChIP-seq datasets.

Indeed, it is important to emphasize that since during cortical neurogenesis, multiple highly coexpressed bHLH factors can bind identical or very similar motifs, certain conclusions drawn from recent epigenomics studies may require some reinterpretation. Several studies highlighted specific proneural factors or E-proteins upon finding enrichment in their associated motifs as described in motif databases such as JASPAR. These enrichments have been interpreted multiple times as evidence of the action of the associated factor, and consequently possibly misled experimental designs for further functional assays, inducing or repressing the expression of these factors^33,45,50^. We propose that motif archetypes as the ones obtained by Vierstra et al.^61^ should be used instead, and complemented, when possible, by the assessment of the correlation between the expression of the transcription factors and the accessibility of the motifs, narrowing down the list of putative candidates, as in Trevino et al. 2021^44^. Moreover, in experimental settings involving TF induction, the silencing of co-expressed TF with similar motif preference may help to disentangle the first order from the secondary order regulatory effects of these TFs. For example, the induction of Neurod1^33^ and *Neurog2*^50^ have shown to induce chromatin remodeling and CpG demethylation, respectively. However, these observations might be primarily explained by the action of other co-expressed factors, *Neurod2* and *Neurod6*. Additionally, a more comprehensive bHLH analysis could clarify the hierarchy of effects on specific targets inferred from knockout studies. For example, Rnd2 was identified as an important target of *Neurog2* on the basis of its expression changes after *Neurog2* knock-out and overexpression in the mouse dorsal telencephalon^13^. However, our data and the study of gene perturbation in the brain of *Neurod2* KO mice suggest that *Neurod2*, acting downstream of *Neurog2*, might actually be the most prominent direct regulator of Rnd2 at E14.5. This is consistent with the observation that *Neurog2* shows reduced ability to induce Rnd2 reporters in E14.5 brain lysates compared with those from earlier stages^26^. This example highlights the importance of assessing simultaneously the action of these highly intermingled proneural factors and E-proteins. Nonetheless, additional ChIP-seq datasets from these other Neurod factors will clarify the level of functional redundancy and cooperation among the members of this family of TFs.

The observed association between NEUROD2 and chromatin remodeling appears to contrast with earlier findings by Fong et al., who reported similar histone H4 acetylation in CAT-CAG and CAG-CAG E-boxes after *Neurod2* induction in the P19 cell line. However, careful examination of their results reveals higher histone acetylation after induction in sites bound by NEUROD2, but not MyoD1, which are those the authors found enriched in CAT-CAG E-boxes. Moreover, while Fong et al. reported a preference for NEUROD2 to bind euchromatin regions^42^, this disparity may be partly attributed to the distinct cell types studied, and it is plausible that the ability of NEUROD2 to bind closed chromatin could depend on the specific molecular context, as exemplified by the case of *Neurog2*, which can reprogram fibroblast to neurons only in the presence of additional factors such as SOX4 or SOX11^82,83^, or Ascl1 which can reprogram pericytes into neurons only when co-expressed with Sox2, another pioneer factor^84^. However, an alternative explanation could be that NEUROD2 exhibits E-box-dependent chromatin remodeling capabilities, as suggested by our analysis. We found that *Neurod2* correlated with the accessibility of CAT-containing E-boxes, however Fong et al. evaluated chromatin accessibility by means of the restriction enzyme PVull^42^, which specifically targets CAGCTG sites, i.e. CAG-CAG E-boxes. This raises the need to re-assess NEUROD2’s chromatin remodeling capabilities, as previously conducted in in vitro induction studies, such as those in Fong et al. 2012^42^ It is conceivable that NEUROD2-bound E-boxes of the CAT-CAT variety may indeed exhibit chromatin remodeling in response to *Neurod2* induction also in this study.

Multiple structural biology studies exploring the connection between TF structure and nucleosome binding coalesce in the principle that pioneer factors are able to bind DNA while avoiding clashes with nucleosomes by using either long kinked or short alpha helices, requiring short or degenerated DNA motifs^85–88^. Since NeuroD factors bind to DNA with long alpha helices that fully contact DNA, Soufi et al. predicted incapacity to bind to closed chromatin^87^. The model also predicts NeuroD factors’ binding to non-degenerated motifs, and in line with this prediction Soufi et al. indicated that NeuroD’s preferred motif is a fixed CAG-CAG E-box with no degenerate positions^87^. However, our observations concerning NEUROD2, adding to previous findings in this and other NeuroD factors^17,42,81^, clearly establishes the CAG-CAG motif as one of many possible E-boxes recognized in vivo and in vitro by NeuroD factors. Moreover, multiple studies inducing ectopic expression of NeuroD, and their closely related NeuroG factors revealed competence to engage closed chromatin, remodel it, and effectively reprogramming various cell types into neurons^42,81,83,89,90^. Finally, a recent study showed that bHLH factors with a CAT half-site binding preference seem to compete more strongly with nucleosomes in vitro than other bHLH factors tested^91^. Although NEUROD2 was not included in this study this represents a potential mechanism to consider. In conclusion, a more nuanced stratification of binding sites based on E-box types might shed new and fresh light into the pioneer capabilities of NEUROD2 and its underlying mechanisms.

## Methods

### ChIP-seq data processing

Raw public NEUROD2 ChIP-seq data of E14.5 NEUROD2, P0 NEUROD2, and E14.5 NEUROG2 was obtained from the Gene Expression Omnibus, with accession numbers GSE67539, GSE63620 and GSE84895, respectively. Fastq reads were aligned to the mouse reference genome mm10 using the bwa-mem program of BWA version 0.7.17^92^ with default parameters. Duplicated reads were removed with SAMtools version 1.12.^93^. We called peaks with MACS3^94^ using uniquely mapped reads, specifying a bandwidth of 200 and using the ChIP-seqs of the GFP of the corresponding replicate as a control. We used only replicates 1 and 2 in both E14.5 and P0 samples, since replicate 3 added almost no additional peaks. For each replicate, peaks were divided into subpeaks using the PeakSplitter program^95^. A final set of consensus peaks was obtained by intersecting subpeaks from the two replicates with the requisite that the summits of both peaks fall within the intersection. NEUROG2 only had one replicate. Venn diagrams of the intersection between NEUROG2 and NEUROD2 were plotted using the *eulerr* R package^96^.

### Single-cell ATAC-seq data processing

We obtained scATAC-seq raw fastq files of the e14.5 mouse cortex experiment from SRA (accession: SRP275808). Reads were aligned to the mouse mm10 assembly using the *count* function of Cell Ranger ATAC version 2.1^97^. We took cell annotations imputed from the scRNA-seq assay of the same study, generously provided to us by Noack et al. We then subdivided the alignment files, taking 350 random cells from each major cell cluster: NSC, IPC, PN1, PN2 and PN3. Bam files from replicates 1 and 2 were merged, and then joined into a single bam file representing the excitatory neuron lineage. Peak calling was performed using the MACS2 program^94^, and the resulting peaks were subsequently divided into subpeaks with PeakSplitter. Subpeaks were centered around the summit and resized to 500bp. Next, taking those peaks and the fragments files of the scATAC-seq assay (GEO accession: GSE155677), we generated count matrices, with the FeatureMatrix function of Signac^98^.

To obtain developmental trajectories of chromatin accessibility, we first ordered cells based on their pseudotime values inputed from the scRNA-seq assay and made groups of 175 cells based on that ordering: 8 groups for NSCs, 5 for IPCs, 7 for PN1, 8 for PN2 and 2 for PN3. Counts from each group of cells were aggregated into pseudobulk samples and normalized as log2 CPMs. We tested peaks for significant differences in accessibility across cell clusters by means of an ANOVA. Peaks with Pvalues>=1e-06 were classified as invariant. To infer trajectories among the remaining dynamic peaks, we first generated predefined combinations of 0s and 1s, representing closed or accessible chromatin, respectively. We generated vectors containing strings of 0s and 1s for each pseudobulk of each cell type and performed Pearson’s correlations of the normalized aggregated accessibilities of the pseudobulks with those vectors. Each peak was assigned to the most highly correlated trajectory. To sort the ATAC-seq peaks based on the correlation with the expression of *Neurog2* and *Neurod2*, we used the Seurat scATAC-seq object with imputed scRNA-seq expressions, from Noack et al.^50^. We sorted cells based on the imputed SCT normalized expression of the transcription factor of interest, performed pseudobulk groups of 50 cells based on that ordering, and correlated the mean SCT imputed expression of the pseudobulk samples with the mean accessibility of each peak.

Open chromatin regions of the P56 mouse cortex were obtained from Li et al.^55^ online resource: http://catlas.org/catlas_downloads/mousebrain/cCREs/. We merged all peaks in one single set. We obtained raw single-cell ATAC-seq matrix from multiple tissues derived from adult mice^56^ from: https://atlas.gs.washington.edu/mouse-atac/data/#release-updates. A peak was considered to be accessible in a given cell cluster if it had at least one count in at least 3% of the cells.

### Gene ontology analysis

To assign genes to NEUROD2 peaks, we made use of the ATAC-seq peak-gene links based on the positive correlation between the accessibility of the peaks and the imputed expression of proximal genes, provided by Noack et al.^50^. A gene was considered linked to a NEUROD2 peak if the NEUROD2 peak intersected with an ATAC-seq peak that positively correlated with the gene.

For the gene ontology analysis of the NEUROD2 peaks in the trajectories, odds ratios were computed as the fraction of genes in each trajectory associated to each ontology term divided by all the genes in that trajectory versus the number of genes associated to each ontology term divided by all the genes used as background by the *g:Profiler*^99^.

### E-box enrichment and centrality

For the motif centrality analysis in the ATAC-seq peaks, .bed files with the occurrences in the mm10 genome of archetypal motifs derived by Vierstra et al.^61^ were obtained from: https://resources.altius.org/#jvierstra/projects/motif-clustering-v2.0beta/. To compute motifs’ centrality in the ATAC-seq peaks, peaks were resized to 1500bp around their summits, and the occurrences of the motifs were counted in two windows: one 50bp around the summit of the peaks and the other 250bp further from the summit. The centrality was computed as the fold change of the occurrences of motifs in the window around the summit vs the distal window, accounting for the lengths in base-pairs of the windows. As a result, motifs that have a centrality value >1 are centrally enriched, and the ones that are <1 are centrally depleted.

### Measurement of CpG methylation levels in E-boxes

BigWig files with whole-genome %CpG methylation for sorted neuronal cells of the embryonic mouse cortex were obtained from Bonev’s lab: https://bonevlab.com/resources. The %CpG methylation data of the excitatory neurons of the adult mouse brain were downloaded as .tsv files from: http://neomorph.salk.edu/hanqingliu/cemba/ALLC/MajorType/, converted to bedGraph files, and then to BigWig files using the *bedGraphToBigWig* program^100^. BigWig files were used as input for the methylation plots. The %CpG methylation heatmaps were plotted using the *EnrichedHeatmap* package^101^, averaging %CpG methylation values in windows of 50bp with the “absolute” method and applying smoothing between the rows of the matrix. The lineplots were produced with the *SeqPlots* package^102^, averaging the %CpG methylation in bins of 10 base-pairs.

### Analysis of HT-selex data

Raw fastq files for each of the four HT-selex rounds performed by Jolma et al.^73^ were downloaded from the European Nucleotide Archive, under accession number ERP001824. For each round, the hexanucleotides were identified by means of regular expressions, and the percentage of sequences that carried each hexanucleotide was computed.

### ModCRE

ModCRE is a structure-based method for predicting transcription factor (TF) binding motifs, represented as position weight matrices (PWMs), and automatically modeling the structure of protein-DNA complexes. It relies on statistical potentials derived from both PDB structures and TF-DNA interactions documented in PBM experiments^103^. Specifically, ES3DCdd statistical potentials, as described in Meseguer et al.^104^, have been found most suitable for predicting PWMs and assessing a TF’s DNA binding capability. The interaction score for a TF-DNA pair is determined by summing the scores of all contact pairs between amino acids and nucleotides within the interaction. To compare different complexes, we employ -ES3DCdd as a measure of binding affinity, where higher values indicate stronger binding. We assess the interaction strength of a specific nucleotide within the binding site by considering all its contacts with interface amino acids, including cases with two or more proteins in the interaction (as seen in dimers and macro-complexes). We applied ModCRE to model the structures of both hetero- and homodimers involving NEUROD2, NEUROG2, and TCF4. We predicted the PWMs and evaluated their binding affinities. To account for structural variability and flexibility, we considered multiple conformations and binding preferences generated from various templates. This approach enhances the robustness of comparisons between homo- and hetero-dimers in our study. The templates used for this analysis were 2QL2, 2YPA, 2YPB, 6OD3, 6OD4, 6OD5, and 1MDY.

### Analysis of E-box sequence conservation

For the genomic variation analysis, we obtained VCF files from the genomes of 154 wild mice^79^. Variants were filtered with GATK^105^, using the following expression: “QD < 2.0 || FS > 60.0 || MQ < 40.0 || MQRankSum < -12.5 || ReadPosRankSum < -8.0”. Then, we removed the indels with VCFtools, and converted the filtered VCF files into bed files. We considered SNVs as polymorphic if they had a Minor Allele Frequency superior or equal to 0.05.

For the evolutionary divergence analysis, we downloaded multiple alignments of the mm10 assembly against 60 species of vertebrates for each chromosome as MAF files, from: http://hgdownload.soe.ucsc.edu/goldenPath/mm10/multiz60way/maf/. The alignments of our motifs of interest were extracted from those MAF files using the *mafsInRegion* tool provided by UCSC (http://hgdownload.cse.ucsc.edu/admin/exe/linux.x86_64/mafsInRegion), converted to fasta files with the maf_to_concat_fasta.py program (https://github.rcac.purdue.edu/kelley/opt-galaxy/blob/master/eggs/bx_python-0.7.2-py2.7-linux-x86_64-ucs4.egg/EGG-INFO/scripts/maf_to_concat_fasta.py), and the fasta files were analyzed using the *read.fasta* function of the *seqinr* package^106^. Only single-nucleotide substitutions were considered, and motifs spanning indels were excluded. For the evolutionary analysis, a motif was considered to be conserved in a given species vs the rat, if it had 0 substitutions in the core CANNTG hexanucleotide. To assign the phylostratigraphical depths to the motifs, we required the motif to be conserved in a proportion of species of a given phylostratigraphical dept:; 20 out of 39 placental mammals, 1 of 3 marsupials, 1 out 1 monotrema and 8 out of 16 non-mammals For the dN/dS analysis, the mm10 vs the rat rn7 genome alignment was downloaded (https://hgdownload.soe.ucsc.edu/goldenPath/mm10/vsRn7/mm10.rn7.synNet.maf.gz), and number of single nucleotide substitutions that occurred within the hexanucleotides were counted, excluding the motifs with indels.

## Acknowledgements

**Funding**: This work was funded by grants MS20/00064 (ISCIII-MICINN/FEDER), PID2019-104700GA-I00 (/AEI/ 10.13039/501100011033), Fundació LaMarató de TV3 and NIH grant R01HG010898-01 (to G.S). BO acknowledges support by PID2020-113203RB-I00 (MCINN, FEDER and AEI DOI: 10.13039/501100011033) and “Unidad de Excelencia María de Maeztu” (ref: CEX2018-000792-M). XdM is supported by the fellowship PRE2020-093064 funded by MCIN/ AEI/10.13039/501100011033.

---

**Extended Data Fig. 1.**
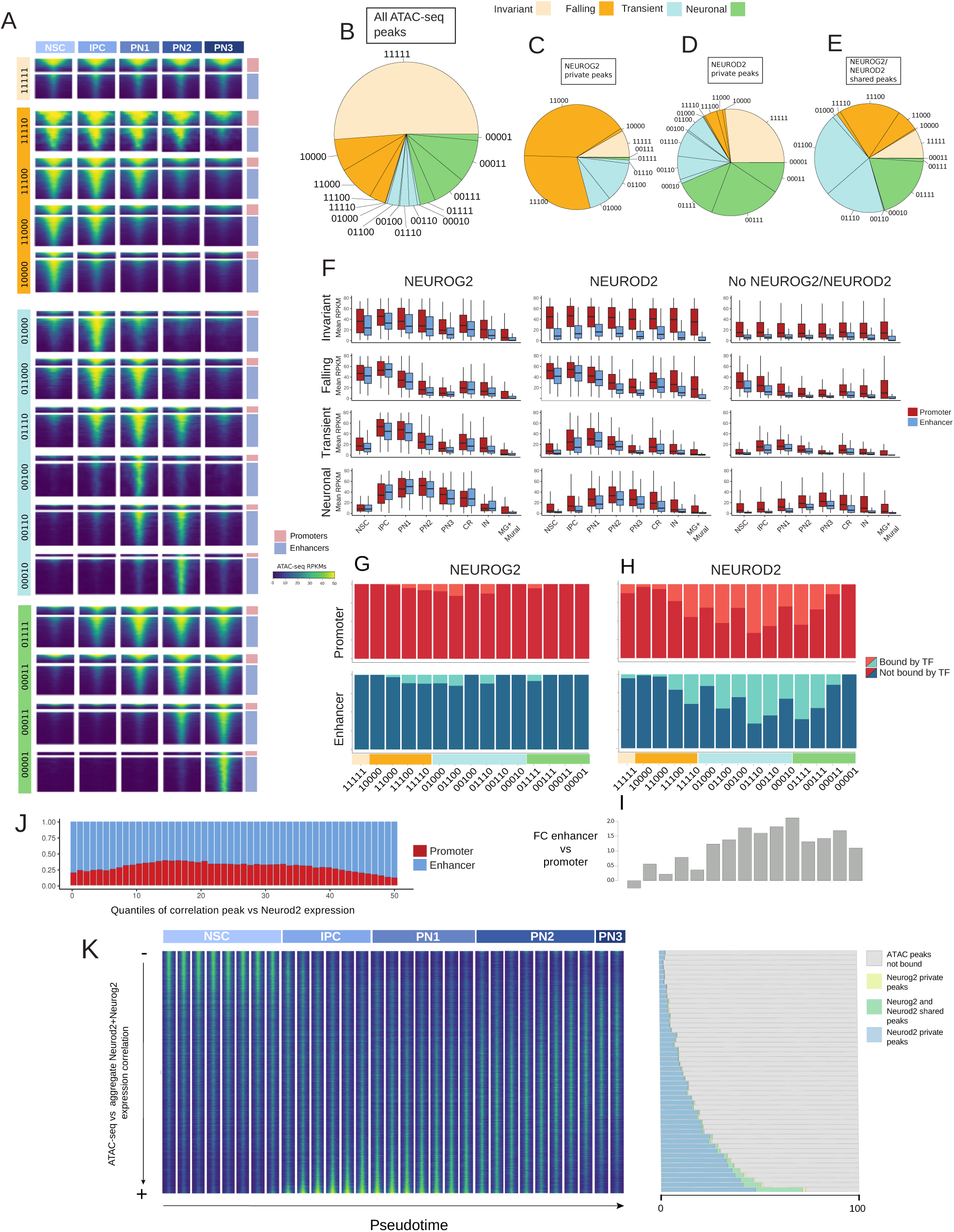
(**A**) Heatmap depicting the ATAC signal found in peaks (rows) classified in pseudotime trajectories (1-accessible, 0-not-accessible chromatin). The proportion of promoter and enhancer regions in each trajectory is shown in the right pie charts. (**B**) Pie chart showing the proportion of pseudotime trajectories among all e14.5 ATAC-seq peaks. (**C**) Pie chart of the proportion of pseudotime trajectories among regions only bound by NEUROG2. (**D**) Pie chart of the proportion of pseudotime trajectories among regions only bound by NEUROD2. (**E**) Pie chart of the proportion of pseudotime trajectories among regions only bound by NEUROG2 and NEUROD2. (**F**) Boxplot showing the distribution of ATAC-seq signals in accessible chromatin peaks of each trajectory overlapping NEUROD2 peaks, NEUROG2 peaks, or none, in each cell type. (**G**) Barplot depicting the proportion of peaks in each trajectory bound by NEUROG2, or (**H**) NEUROD2. (**I**) Barplot indicating the fold change of the proportion of NEUROD2-bound peaks in each trajectory in promoters versus enhancers. (**J**) Barplot representing the proportion of enhancers and promoters in quantiles of ATAC-*Neurod2* correlation. (**K**) (Left) Heatmap representing pseudobulk signal around the summits of the ATAC-seq peaks sorted by the correlation of their accessibility with the combined expression of NEUROG2 and *Neurod2*. Pseudobulk samples are composed of cells grouped in increasing pseudotime bins. (Right) Proportion of ATAC-seq peaks bound by NEUROG2 and NEUROD2 in each of 50 quantiles of the correlation with the corresponding transcription factor.

**Extended Data Fig. 2.**
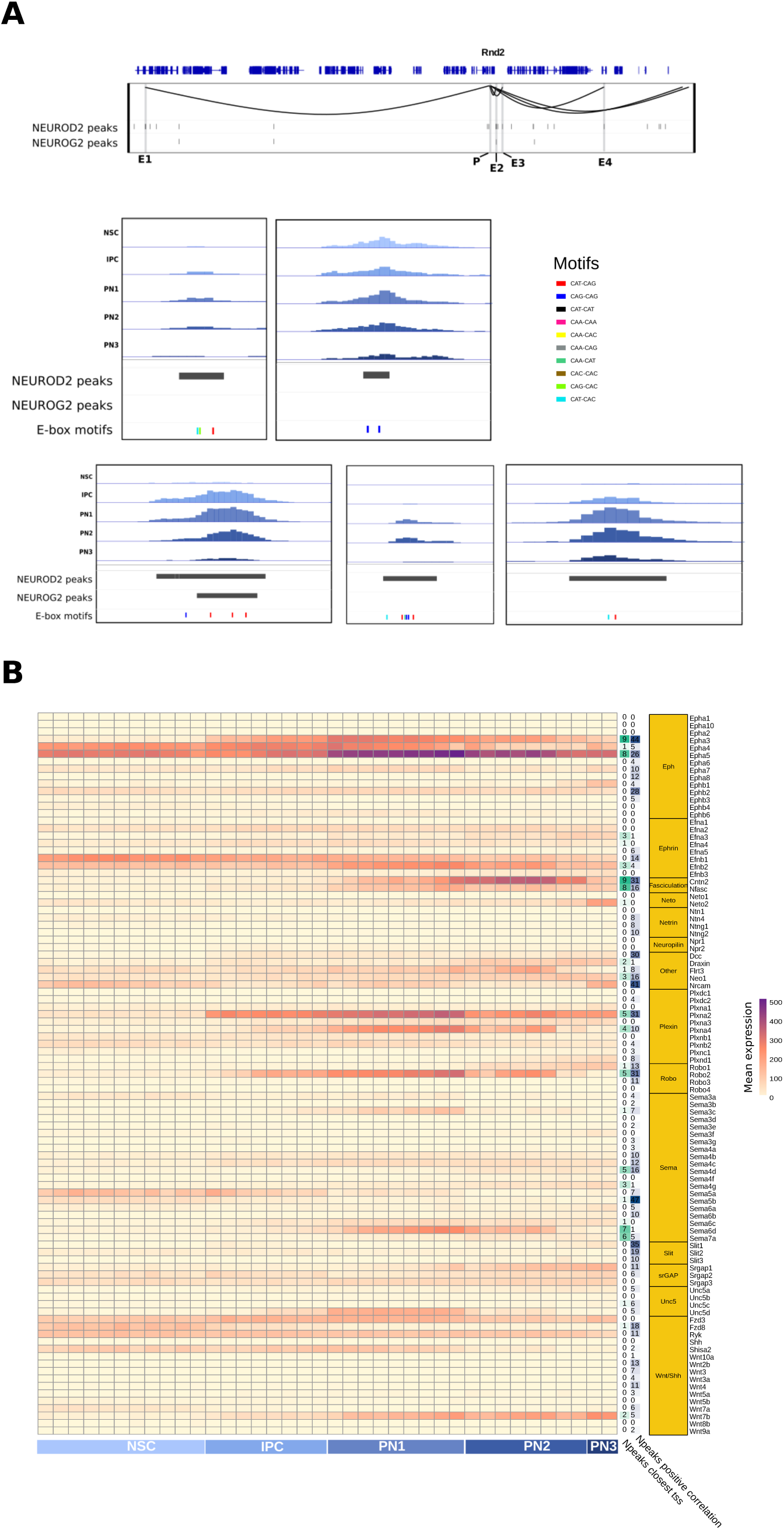
(**A**) Genome browser representation of NEUROD2 and NEUROG2 peaks associated with the gene *Rnd2* and their signal. (**B**) Heatmap representing average expression of axon guidance genes on pseudobulk samples ordered by pseudotime. Right columns contain the number of NEUROD2 peaks associated with each axon guidance gene (right), and the number of NEUROD2 peaks correlated with target gene expression (left).

**Extended Data Fig. 3.**
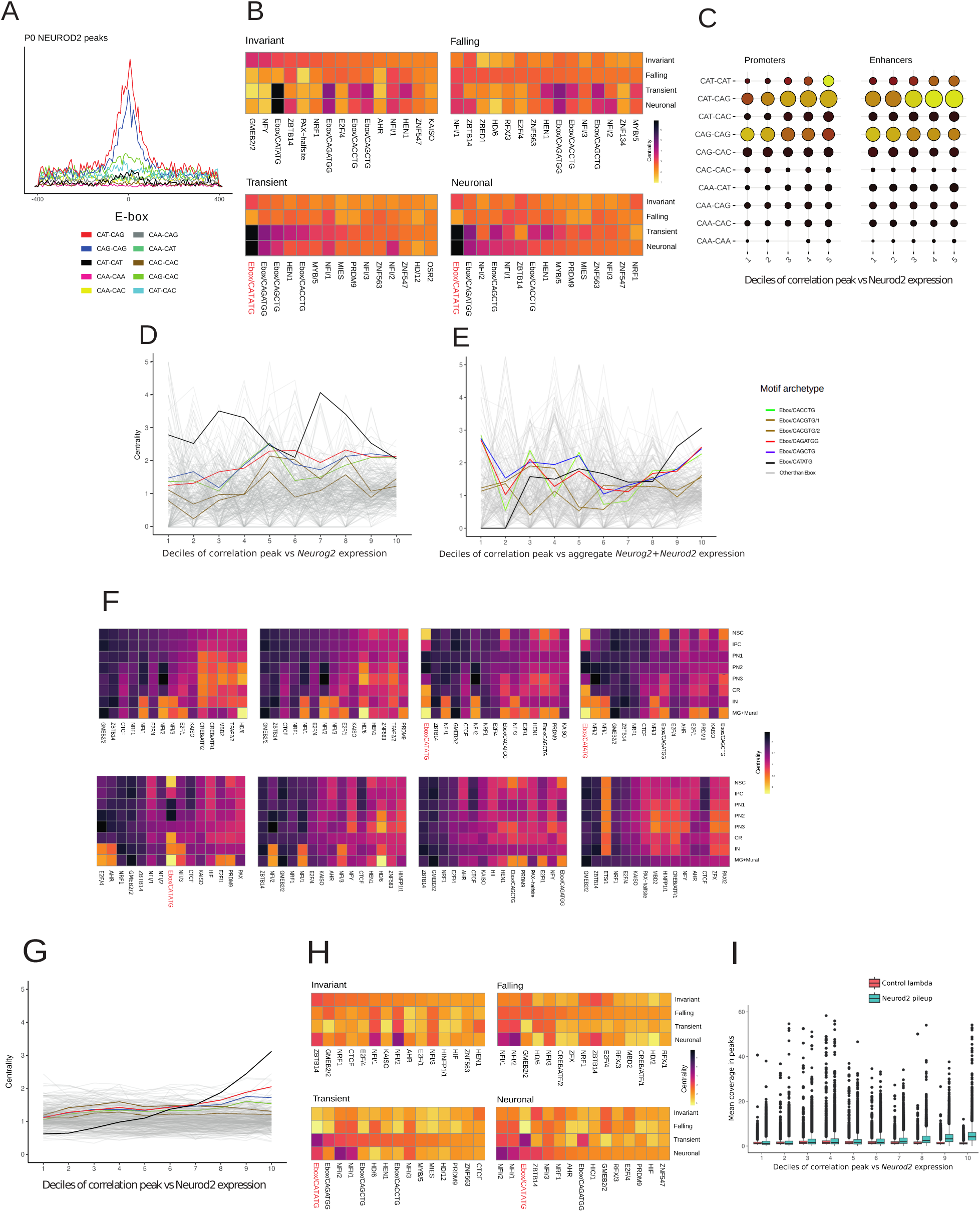
(**A**) Number of E-box motifs per base pair and per peak around the summits of NEUROD2 peaks in P0 mouse brain. (**B**) Heatmap representing centrality enrichment for 233 archetype motifs in ATAC peak bound by NEUROD2 across the four major trajectories. Top 15 are shown. (**C**) Centrality (motifs 50bp around the summit vs >150bp further) of each E-box type around the summits of the NEUROD2 peaks overlapping promoters and enhancers, binned by the correlation with *Neurod2* expression levels. (**D**) Line plot indicating motif centrality (motifs 50bp around the summit vs >250bp further) of all the archetypal motifs determined in Vierstra et al. around the summits of the ATAC-seq peaks, grouped by the correlation of their ATAC accessibility with the expression levels of NEUROG2. (**E**) Line plot indicating motif centrality (motifs 50bp around the summit vs >250bp further) of all the archetypal motifs determined in Vierstra et al.^61^ around the summits of the ATAC-seq peaks not bound by NEUROD2 and NEUROG2, grouped by the correlation of their ATAC accessibility with the aggregated expression levels of *Neurod2* and *Neurog2*. (**F**) Heatmap representing centrality enrichment for 233 archetype motifs across ATAC-seq peaks derived from all cell types in the E14.5 mouse cortex. (**G**) Line plot indicating motif centrality (motifs 50bp around the summit vs >250bp further) of all the archetypal motifs determined in Vierstra et al.^61^ around the summits of the ATAC-seq peaks not bound by NEUROD2, grouped by the correlation of their ATAC accessibility with the expression levels of *Neurod2*. (**H**) Heatmap representing centrality enrichment for 233 archetype motifs across the four major ATAC peak trajectories not bound by NEUROD2 or NEUROG2. Top 15 are shown. (**I**) Boxplot depicting NEUROD2 ChIP-seq signal in ATAC-peaks not bound by NEUROD2 as a function of the correlation of their ATAC accessibility with the expression levels of *Neurod2*. A similar analysis using background genomic regions is shown for comparison.

**Extended Data Fig. 4.**
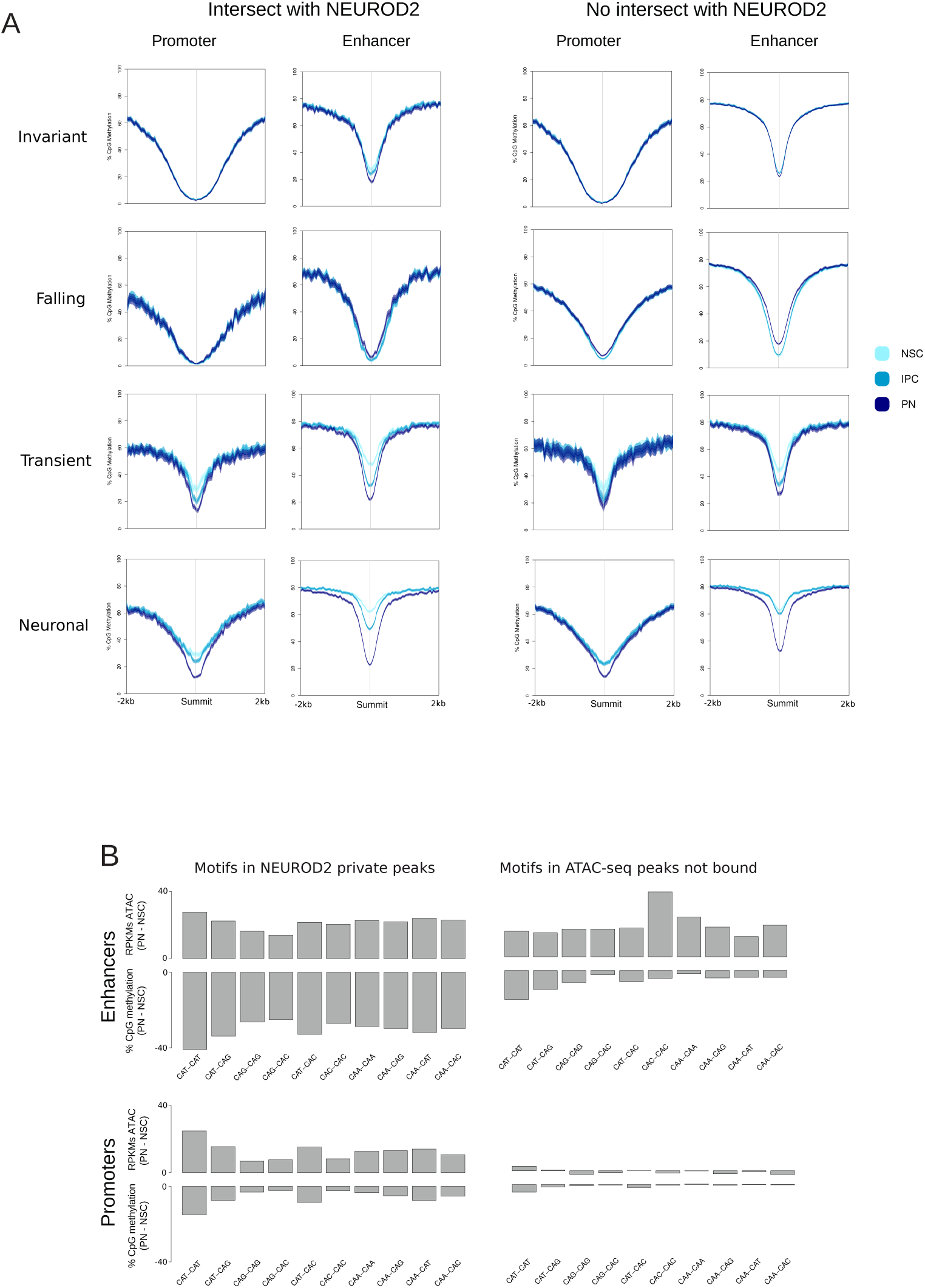
(**A**) Lineplot representing the average signal of the subsets of ATAC peaks in each trajectory bound or not bound by NEUROD2, with ribbons representing the standard error of the mean. (B) Barplots representing the difference in percentage of CpG methylation measured 5bp around each type of E-box between PN and NSC (bottom) in promoters and enhancers bound by NEUROD2. The top barplots represent the same comparisons using ATAC-seq RPKMs.

**Extended Data Fig. 5.**
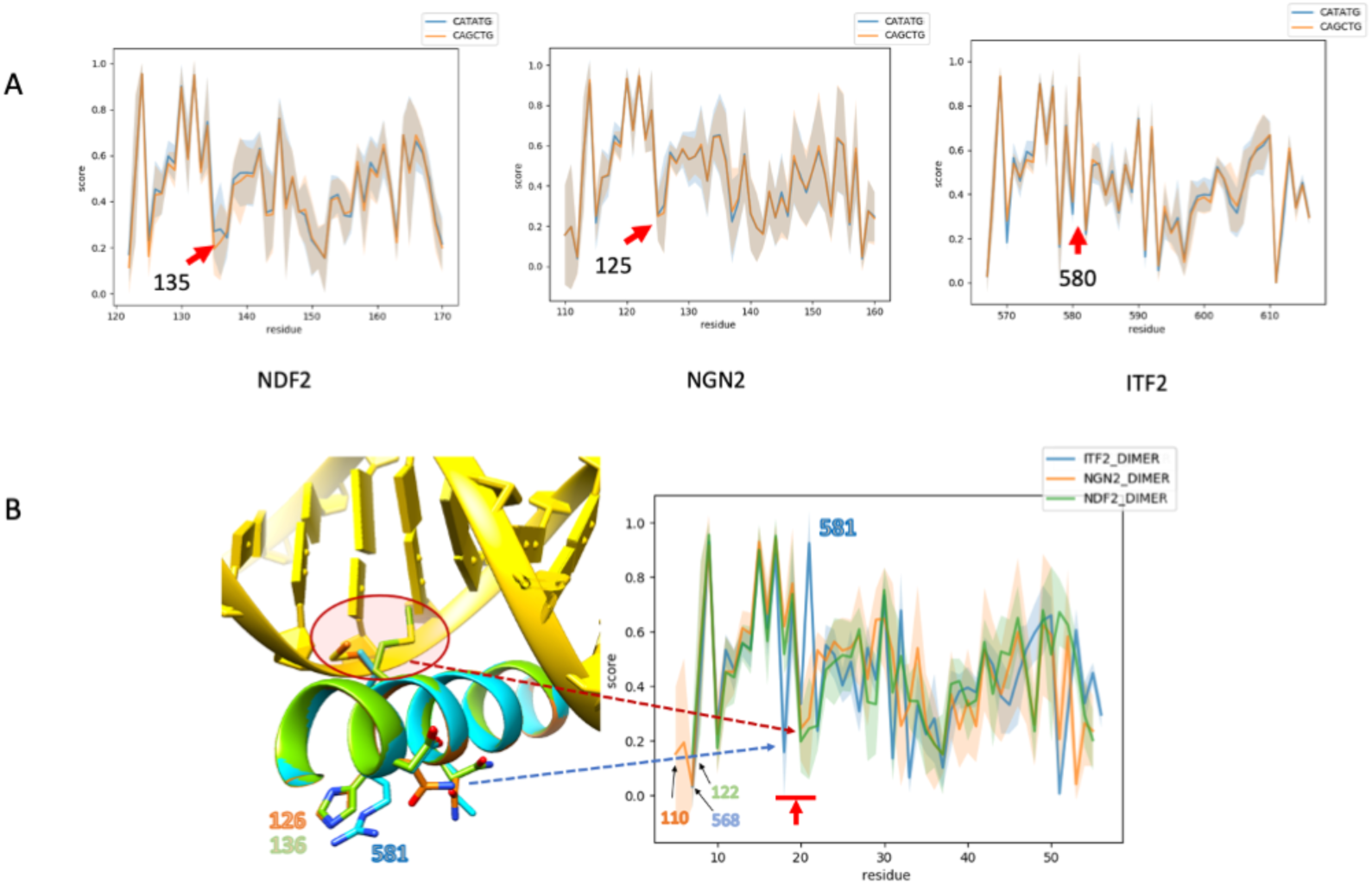
(**A**) The figures depict normalized interaction scores between DNA sites and amino acids of NEUROD2, NEUROG2, and TCF4 for CAT-CAT (blue) and CAG-CAG (orange) sequences. Red arrows highlight score differences and their position in the amino-acid sequence. (**B**) The plot at the right displays normalized scores per amino acid for NEUROD2 (green), NEUROG2 (orange), and TCF4 (blue) homodimers. On the left, a ribbon-plot illustrates the modeled bound helix of NEUROD2 (green), NEUROG2 (orange), and TCF4 (blue), revealing encircled in red side-chains of Val-580 in TCF4 and Met-135/Met-125 in NEUROD2/NEUROG2. The visuals provide insights into sequence-specific interactions and structural distinctions among the analyzed transcription factors. Side-chains of Arg-581 in TCF4 and His-136/His-125 in NEUROD2/NEUROG2 provide reference points in the plot, with reassigned numbers for correct alignment.

**Extended Data Fig. 6.**
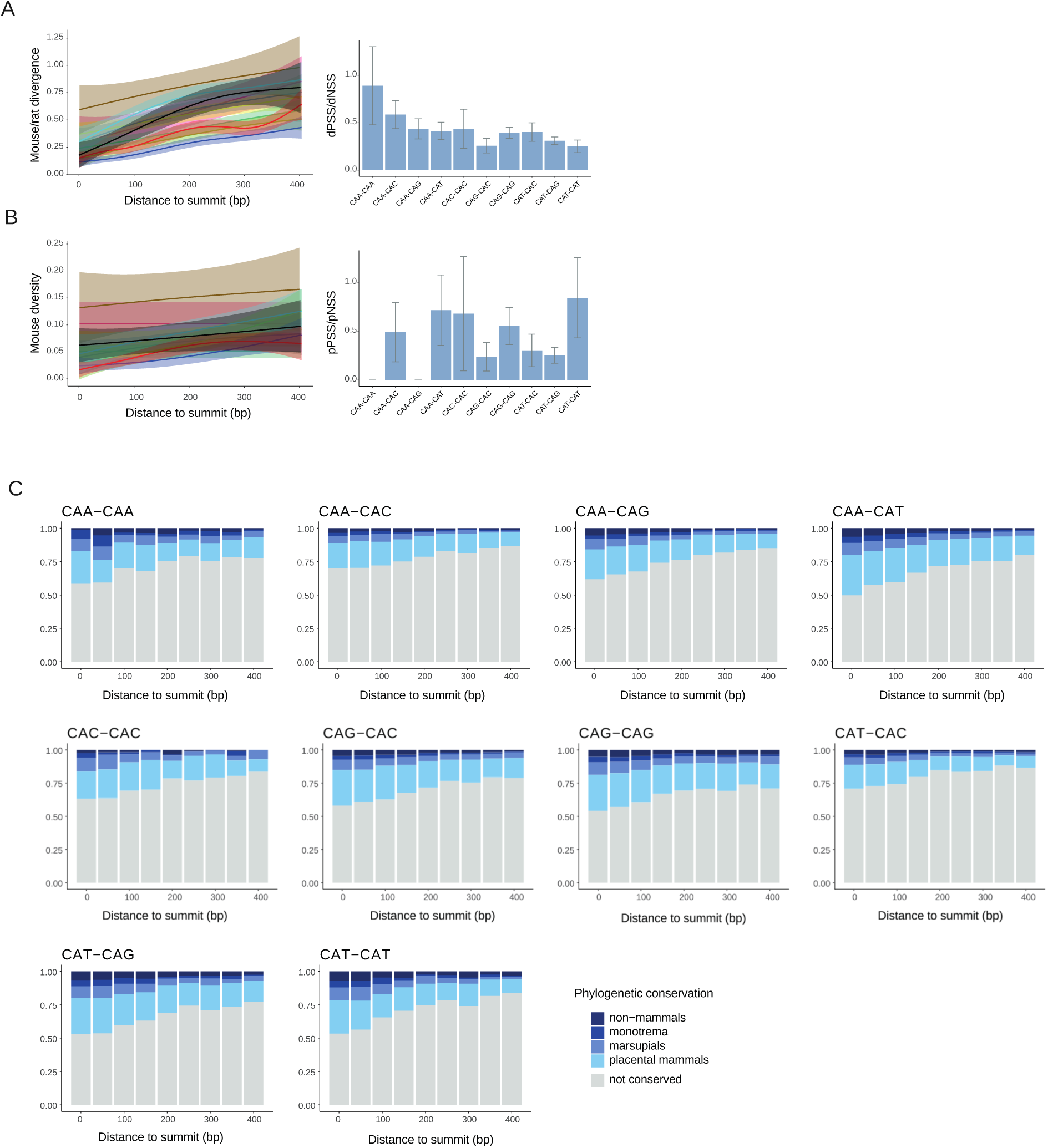
(**A**) Line plots displaying the number of mouse-rat substitutions within E-boxes in bins of increasing distance to NEUROG2 peak summits (left). Barplot showing the ratio of the dPSS and dNSS in each E-box type in NEUROD2 peaks (right). (**B**) Line plots showing number of SNPs within E-boxes in bins of increasing distance to NEUROG2 peak summit (left). Barplot showing the ratio of the pPSS and pNSS in each E-box type in NEUROD2 peaks (right). (**C**) Barplots representing the proportion of each type of E-boxes displaying various degrees of phylogenetic conservation at bins of increasing distance to NEUROD2 peak summits.

**Extended data figure 7.**
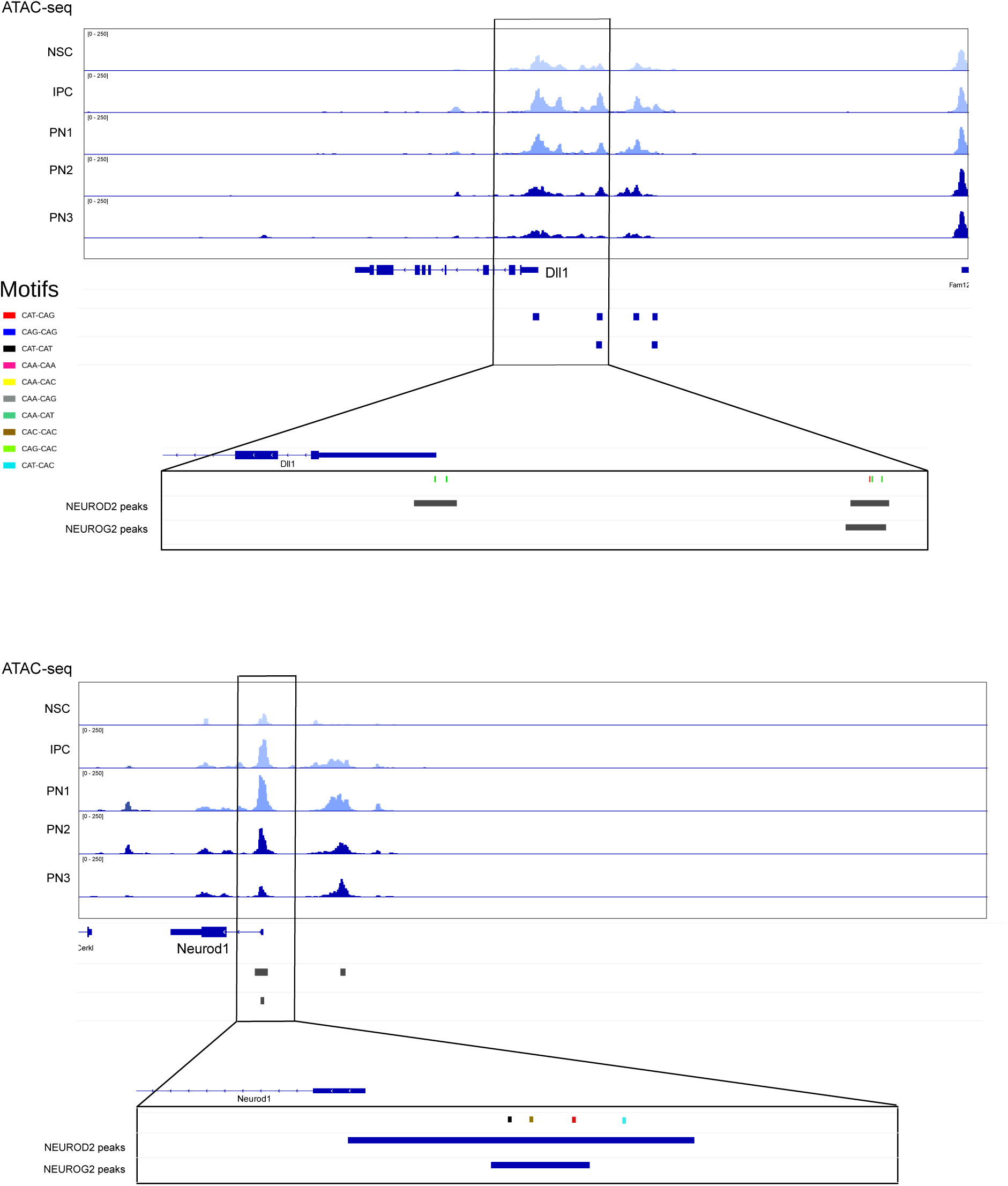
Genome browser representation of NEUROD2 and NEUROG2 peaks associated with the gene *Dll1* and *Neurod1* and their signal. Specific E-boxes overlapping highlighted peaks are represented.

